# Nitric Oxide modulates spontaneous Ca^2+^ release and ventricular arrhythmias during β-adrenergic signalling through *S*-nitrosylation of Calcium/Calmodulin dependent kinase II

**DOI:** 10.1101/2023.08.23.554546

**Authors:** Amelia S. Power, Esther Asamudo, Luke P. I. Worthington, Chidera C. Alim, Raquel Parackal, Rachel S. Wallace, Obialunanma V. Ebenebe, Joan Heller Brown, Mark J. Kohr, Donald M. Bers, Jeffrey R. Erickson

**Affiliations:** Department of Physiology and HeartOtago, University of Otago, Dunedin, New Zealand; Department of Physiology, University of Auckland, Auckland, New Zealand; Department of Pharmacology, University of California, Davis; Department of Environmental Health and Engineering, Johns Hopkins Bloomberg School of Public Health, Baltimore, Maryland; Department of Pharmacology, University of California, San Diego, La Jolla

**Keywords:** Arrhythmias, S-nitrosylation, CaMKII, Calcium

## Abstract

**Rationale:** Nitric oxide (NO) has been identified as a signalling molecule generated during β-adrenergic receptor (AR) stimulation in the heart. Furthermore, a role for NO in triggering spontaneous Ca^2+^ release via *S*-nitrosylation of Ca^2+^/calmodulin kinase II delta (CaMKIIδ) is emerging. NO donors are routinely used clinically for their cardioprotective effects in the heart, but it is unknown how NO donors modulate the pro-arrhythmic CaMKII to alter cardiac arrhythmia incidence.

**Objective:** We test the role of *S*-nitrosylation of CaMKIIδ at the Cys-273 inhibitory site and Cys-290 activating site in cardiac Ca^2+^ handling and arrhythmogenesis before and during β-AR stimulation.

**Methods and Results:** We measured Ca^2+^-handling in isolated cardiomyocytes from C57BL/6J wild-type (WT) mice and mice lacking CaMKIIδ expression (CaMKIIδ-KO) or with deletion of the *S*-nitrosylation site on CaMKIIδ at Cys-273 or Cys-290 (CaMKIIδ-C273S and -C290A knock-in mice). Cardiomyocytes were exposed to NO donors, S-nitrosoglutathione (GSNO; 150 μM), sodium nitroprusside (SNP; 200 μM) and/or β-adrenergic agonist isoproterenol (ISO; 100 nM). WT and CaMKIIδ-KO cardiomyocytes treated with GSNO showed no change in Ca^2+^ transient or spark properties under baseline conditions (0.5 Hz stimulation frequency). Both WT and CaMKIIδ-KO cardiomyocytes responded to ISO with a full inotropic and lusitropic Ca^2+^ transient response as well as increased Ca^2+^ spark frequency. However, the increase in Ca^2+^ spark frequency was significantly attenuated in CaMKIIδ-KO cardiomyocytes. The protection from ISO-induced Ca^2+^ sparks and waves was mimicked by GSNO pre-treatment in WT cardiomyocytes, but lost in CaMKIIδ-C273S cardiomyocytes that displayed a robust increase in Ca^2+^ waves. This observation is consistent with CaMKIIδ-C273 *S*-nitrosylation being critical in limiting ISO-induced arrhythmogenic sarcoplasmic reticulum Ca^2+^ leak. When GSNO was applied after ISO this protection was not observed in WT or CaMKIIδ-C273S but was apparent in CaMKIIδ-C290A. In Langendorff-perfused isolated hearts, GSNO pre-treatment limited ISO-induced arrhythmias in WT but not CaMKIIδ-C273S hearts, while GSNO exposure after ISO sustained or exacerbated arrhythmic events.

**Conclusions:** We conclude that prior *S*-nitrosylation of CaMKIIδ at Cys-273 can limit subsequent β-AR induced arrhythmias, but that *S*-nitrosylation at Cys-290 might worsen or sustain β-AR-induced arrhythmias. This has important implications for the administration of NO donors in the clinical setting.

## Introduction

The ability of the heart to rapidly enhance output is mediated in part by stimulation of β-adrenergic receptors (β-AR), which trigger increased release of intracellular Ca^2+^ from the sarcoplasmic reticulum (SR) and accelerated re-uptake within cardiomyocytes ^1,2^. Excessive β-AR stimulation can lead to arrhythmias ^3^ and heart failure ^4^ therefore, an understanding of the down-stream signalling pathways that lead to pathological Ca^2+^ handling is vital. Nitric oxide (NO) is a gaseous signalling molecule that can alter cardiac function ^5^ and is produced endogenously within cardiomyocytes following β-AR stimulation ^6^. NO-releasing drugs (donors) have been used clinically for over a century for their protective actions on heart function ^7^ however, the direct effects of NO on cardiomyocytes is still under investigation. NO can exert positive inotropic effects due to protein modification by *S*-nitrosylation ^8^, where NO covalently attaches to cysteine (Cys) residues and alters protein activity ^9^. NO exposure (either endogenously produced or exogenously applied) has been regarded as cardioprotective in the context of ischemia-reperfusion, due to the *S*-nitrosylation of cardiac proteins ^10^.

Recently, evidence has demonstrated that endogenous NO production is linked to increased spontaneous release of Ca^2+^ from the SR during β-adrenergic stimulation ^11–13^. These observations challenge the cardioprotective role of NO, as spontaneous Ca^2+^ leak is arrhythmogenic ^14^. The function of several cardiac Ca^2+^ handling proteins is reported to be modulated by *S*-nitrosylation, including the ryanodine receptor type 2 (RyR2), L-type Ca^2+^ channel (LTCC) and SR Ca^2+^ ATPase (SERCA) ^15^. Interestingly, NO can alter the frequency of Ca^2+^ sparks in cardiomyocytes following β-AR stimulation in either a positive or negative manner ^16^, although the detailed mechanism by which NO can both enhance and reduce Ca^2+^ spark frequency is not clearly understood.

An emerging target for cardiac NO is the Ca^2+^/calmodulin (CaM) dependent kinase II delta (CaMKIIδ) ^11,13,17^, a nodal regulator of cardiac Ca^2+^ handling ^18^. The regulatory domain of CaMKIIδ contains two *S*-nitrosylation sites that alter its activity ^19^. *S*-nitrosylation at Cys-290, following initial activation by Ca^2+^/CaM, causes autonomous activation of the kinase, consistent with the observation that NO exposure can enhance CaMKIIδ activity and increase Ca^2+^ sparks ^11,13,20^. In contrast, *S*-nitrosylation at Cys-273 inhibits CaMKIIδ by preventing activation by Ca^2+^/CaM ^19^, suggesting a dual role for NO in mediating CaMKIIδ activity that may be alternately protective or pathological depending on intracellular conditions.

β-AR stimulation increases spontaneous Ca^2+^ release from RyR2 in a CaMKIIδ-dependent manner^4^. β-AR stimulation also increases endogenous NO production in cardiomyocytes ^13^, which is necessary for the enhancement of Ca^2+^ sparks ^11^. We therefore hypothesized that arrhythmogenic activity at the cellular and whole heart levels would be prolonged by NO after β-adrenergic stimulation due to *S*-nitrosylation of the Cys-290 site on CaMKIIδ. Further, we hypothesized that exposure to NO before β-AR stimulation would lead to *S*-nitrosylation of the Cys-273 site on CaMKIIδ, reducing β-AR induced Ca^2+^ mishandling. Here, we tested the role of NO and CaMKIIδ on altered Ca^2+^ handling in the context of β-AR stimulation in isolated mouse cardiomyocytes and Langendorff-perfused mouse hearts from transgenic mice lacking expression of CaMKIIδ or the *S*-nitrosylation sites at Cys-273 (inhibitory) or Cys-290 (activating).

## Materials and Methods

### Mouse models

Experiments were performed using 12 - 16-week old male and female mice with four genotypes: C57BL/6J wild-type (WT), knockout mice with deletion of CaMKIIδ (CaMKIIδ-KO), and novel knock-in mouse models with a single mutation of the Cys-273 or Cys-290 *S*-nitrosylation sites on CaMKIIδ (CaMKIIδ-C273S and CaMKIIδ-C290A). The CaMKIIδ-C290A knock-in mice were generated by the UC Davis Mouse Biology Core and have been described previously ^20^. CaMKIIδ-C273S animals were generated using CRISPR/Cas9 genome editing at the Australian Phenomics Facility (Australian National University, Australia). The cysteine codon “TGT” at position 273 was replaced with a serine codon “TCT” using two single guide RNAs: AGTGACTTAC**ACA**GATCCAT**GGG** and **CCC**ATGGATC**TGT**GTAAGTCACT and an oligonucleotide repair template: TTCTGTAAGATTATTTTAAACTTATGAAAAGTGACTAGG GGTTTATCCTTCAGTTTGGCTCCTGGGGTGCATGCGAA<u>AGTGACTTAC**AGA**GATCCAT</u>**GGG**T GTTTCAGGGCCTCAGAGGCTGTGATACGTTTGGCAGGGTTGATGGTCAGCATTTTGTTGATGAGGTCTTTGG. All genetically modified animals were crossed with C57BL/6J mice to share common genetic background. CaMKIIδ-C273S mice were genotyped via PCR amplification of the oligonucleotide repair template with the following primers ATCTCATGTATACCAGGTTTCCC and CCTTCTGGGGTGCAGTAAGT and then ABI BigDye Terminator sequencing of that product. Ear notches from mice were incubated in 100µl of lysis buffer (100 mM Tris-HCL, 5 mM EDTA, 200 mM NaCl, 0.2% SDS, pH 8.5 with Proteinase K at a final concentration of 100 ug/mL) for 3 hrs at 55°C, 15 min at 95°C and then stored overnight at 4°C. 1 µl of the lysate was used for template in 20 µl reactions. PCR was carried out using Phusion High-Fidelity DNA Polymerase (Thermo Fisher) with primers at 0.5 µM and initial denaturation of 1 min at 98°C; 35 cycles of 98°C for 10 sec, 62°C for 15 seconds (s), 72°C for 15 s; and the final extension at 72°C for 10 min. Products were run on 1% agarose gels and then gel-purified using the NucleoSpin Gel and PCR Clean-up kit (Macherey-Nagel) according to manufacturer’s instructions. This was then submitted for sequencing to Genetic Analysis Services at University of Otago with either of the primers listed above. Results were analyzed with SnapGene Viewer (Dotmatics).

Mice were fed Teklad Global 18% Protein Rodent diet (Envigo) *ad libitum* and housed at 20-22°C under 12-hour light-dark cycles. The Animal Ethics Committee at both the University of Otago (AUP-18-42) and University of California, Davis (21572) approved use of animals in this study.

### Protein expression

The ventricles were dissected and snap frozen in liquid nitrogen for protein analysis as previously described^21^. Briefly, approximately 10 – 20 mg of ventricle tissue was homogenized in 20 volumes of buffer containing (in mM) 50 Tris-HCl pH 7.4 150 NaCl, 1 EDTA, 1 phenylmethylsulfonyl fluoride, 0.1% SDS, 1% triton X-100 (CaMKIIδ -KO) or 50 Tris-HCl pH 7.5, 1 phenylmethylsulfonyl fluoride, 3% SDS (RyR, SERCA, CaMKIIδ-C273S) and supplemented with cOmplete protease inhibitor (Roche). Lysates were then incubated on ice for 15 min and centrifuged for 15 min at 15000 rcf at 4°C. Lysates were stored at −80°C until use.

For CaMKIIδ expression in the CaMKIIδ-KO mice, 25μg of protein homogenate was separated on 10 % SDS polyacrylamide gel and transferred for 3 hrs on ice at 100 V to PVDF membrane, and then blocked for 1 hr at room temperature (RT) in 5% non-fat milk powder in Tris-bufferedsaline, 0.05% Twee-20 (TBST). CaMKIIδ was probed for using a primary antibody against CaMKIIδ (1:5000, ThermoFisher PA5-22168) and GAPDH (1:10000, GeneTex GTX627408) for a loading control. Blots were then incubated with secondary mouse or rabbit antibodies conjugated with horse-radish peroxidase (1:10000; Pierce 31460 and 31430), visualized by chemiluminescence detection with Super-signal West Pico (Thermo Fisher), and imaged using a Syngene gel doc system.

For the CaMKIIδ-C273S mice, 5 μg of protein homogenate was separated on Criterion™ XT Bis-Tris (RyR, SERCA, Actin) or 4–15% Mini-PROTEAN® TGX™ Precast stain-free (CaMKIIδ) gels (Bio-rad) and transferred at 4°C for 16 hrs at 30 V onto 0.45 µm supported nitrocellulose membrane (Bio-rad). Membranes were blocked in EveryBlot Blocking Buffer (Bio-rad) for 20 min at RT. Primary antibodies for RyR [34C] (1:500, Abcam ab2868), SERCA2a (1:5000, Badrilla A010-20), and CaMKIIδ (1:2000, Thermo Fisher PA5-22168) were incubated in EveryBlot Blocking Buffer (RyR) or TBST for 2 hrs at RT followed by mouse (1:20000, Pierce 31430) or rabbit (1:10000, Pierce 31460) secondary antibodies conjugated with horse-radish peroxidase in TBST for 1 hr at RT. Anti-beta Actin [AC-15] antibody HRP (1;20000, Abcam ab49900) was incubated in TBST for 1 hr at RT. Blots were visualized by chemiluminescence detection with Clarity Western ECL Substrate (Bio-rad) and imaged using a ChemiDoc MP (Bio-rad).

### Heart Fibrosis

When ventricles were dissected as noted above the apex of the heart was fixed in 4% formalin (Sigma) for 24 hrs. The fixed tissue was then transferred to Phosphate-buffered saline (PBS) for 24 hrs then 30% sucrose in PBS for 48 hrs. The samples were then rinsed in PBS and cryopreserved in FSC 22 Frozen Section Media (Leica) and stored at −80°C until use. Sections were cut (8 µm) in a Leica CM 1950 Cryostat and 4 sections/sample mounted onto Superfrost Plus microscope slides (LabServ). Slides were stored at −20°C for up to one month before staining. Masson’s trichrome staining of sections was performed by the Histology Unit at University of Otago. Slides were then scanned on an Aperio Slide Scanner (Leica) under 40X magnification.

Using Leica’s ImageScope software, randomly selected regions of each section from blinded files were extracted and saved as tiff files. Files were imported into Adobe Photoshop and the background removed. After this, total pixels of the region were determined. Blue target pixels (representing collagen) were then identified. Percent collagen is the sum of all regions from sample sections of target pixels divided by total pixels.

### Measurement of NO concentration

The amount of NO released following addition of NO donor S-nitrosoglutathione (GSNO) to our experimental buffer (KRH; see below) was measured using an Apollo 1000 Free Radical analyser with an ISO-NOPF100 NO microsensor (1 mm) (World Precision Instruments, USA). The sensor was calibrated with S-nitroso-N-Acetyl-D,L-Penicillamine (Toronto Research) in 0.1 M CuCl_2_ solution ^22^. Data were acquired using a Powerlab 2/25 and recorded in LabChart 8.1 (ADInstruments, New Zealand).

### Isolation of cardiomyocytes

Ventricular cardiomyocytes were isolated from mouse hearts using a Langendorff-perfusion method ^23^. Mice were anesthetized with isoflurane until loss of toe pinch reflex. Hearts were excised, aorta cannulated and subjected to retrograde perfusion with either Ca^2+^-free Krebs-Ringer HEPES (KRH) Buffer containing (in mM): 125 NaCl, 5 KCl, 1.2 MgCl_2_, 25 HEPES, 6 glucose, pH 7.4 at 37°C or MEM solution (for CaMKIIδ-C290A mice at UC Davis the KRH for myocyte isolation containing (in mM): 135 NaCl, 4.7 KCl, 1.2 MgSO_4_•7H_2_O, 0.6 KH_2_PO_4_, 0.6 Na_2_HPO_4_•7H_2_O, 10 glucose, 20 HEPES, pH 7.4) gassed with 100% O_2_. Hearts were then perfused for 5 - 10 min with 0.125 mM CaCl_2_ KRH buffer supplemented with 1mg/mL of type II collagenase (Worthington Biochemical). Ventricles were cut into 0.1 mM CaCl_2_ solution, and the tissue was minced into small pieces and further dissociated with gentle pipetting. Cardiomyocytes were collected by filtering solution though a 200 μm mesh and CaCl_2_ concentration gradually increased.

### Calcium imaging

Freshly isolated cardiomyocytes were loaded with 2 μM Fluo-4-AM (Thermo Fisher) for 20 min at room temperature, followed by wash and de-esterification for 30 min. Cells were added to a stimulation bath mounted on either a Nikon A1+ confocal laser scanning microscope (excited at 488 nM and emission as collected through a 525/50 nm filter using a 63X oil immersion objective) or Bio-Rad Radiance 2100 (40X oil immersion objective lens) for CaMKIIδ-C290A myocytes only. Cardiomyocytes were field stimulated at 0.5 Hz for 30 s to establish a steady-state prior to recording Ca^2+^ transients in line-scan mode (2 ms/line, 0.15 x 0.15 μm pixel size). Ca^2+^ sparks and the occurrence of waves were measured under quiescent conditions (30 s after termination of 0.5 Hz pacing). Total SR Ca^2+^ content was determined at the end of each experiment with a rapid 20 mM caffeine exposure in Ca^2+^-free KRH following a 30 s train of 0.5 Hz stimulations. Steady-state Ca^2+^ transients were averaged, and parameters analysed in custom written MATLAB software (version R2018a, MathWorks) and custom code written in Python. Ca^2+^ sparks were detected using the SparkMaster plugin in Fiji using a detection criteria of 3.8 ^24^. All experiments were performed at room temperature and cardiomyocytes were constantly perfused with KRH buffer containing 1.5 mM CaCl_2_ unless stated otherwise. Cardiomyocyte dimensions were measured in Fiji from confocal images acquired using frame-scan mode.

### Animal Echocardiography

Echocardiography was carried out on WT and CaMKIIδ-C273S at 12 weeks of age using a Vivid E9 Imaging System (General Electric Vingmed Ultrasound, Norway) equipped with an ultrasound probe of 11 MHz frequency. Animals were maintained under anaesthesia with 0.5 L/min of 100% oxygen, 1-2% isoflurane. All captures were obtained and analysed as described previously ^21^.

### Isolated Heart Function

Under anaesthesia (3% isoflurane, 0.5 L/min of 100% O_2_), mice were injected intraperitoneally with heparin (10,000 U/kg). After 5 min, isoflurane inhalation was further increased to 5% to induce deep anaesthesia. The hearts were excised and arrested in Ca^2+^-free Krebs-Henseleit buffer (KHB) on ice before being cannulated via the aorta and Langendorff-perfused with modified KHB. The modified KHB solution contained (in mM): 2.0 CaCl_2_.2H_2_O, 118.5 NaCl, 4.7 KCl, 1.2 MgO_4_S.7H_2_O, 1.2 KH_2_PO_4_.H_2_O, 25 NaHCO_3_, 11 glucose and 2.0 pyruvate. The modified KHB was aerated continuously with 95% oxygen and 5% carbon dioxide to maintain pH of 7.4. The coronary pressure was maintained at 75 mmHg and coronary flow rate range was between 1.5 - 4 mL/min. Each heart was allowed to stabilize in the organ bath while continuously perfused with KHB at 37°C for 5 min. Afterwards, the left atrium was removed and a balloon connected to a pressure transducer (ADInstruments) was inserted into the left ventricle (LV) via the atrial appendage. The balloon was inflated with distilled water to achieve a Left Ventricular End Diastolic Pressure (LVEDP) of 5 - 10 mmHg. This was followed by a stabilization period of 30 min, during which the LV pressure trace was monitored for regular beats before obtaining basal heart function parameters.

Baseline data were recorded for 10 min, followed by drug infusion for 10 min. To determine the effect of NO and β-AR stimulation on heart function, the mouse hearts were randomly assigned to three groups depending on the order of drug infusion. The control group received isoproterenol (ISO, 100 nM) only followed by wash out with control KHB buffer. The treatment groups received either ISO (100 nM) followed by S-nitrosoglutathione (GSNO, 150 µM) or GSNO (150 µM) before ISO (100 nM). LV pressure, coronary flow, and temperature were recorded using a data acquisition system (LabChart 8.1, ADInstruments). The trace obtained was saved for offline analysis to measure the incidence of arrhythmias. Ventricular arrhythmic events from the LV pressure trace were evaluated and classified as outlined in Supplementary Figure 2 and Table 1. The different types of arrhythmias observed at least once in the isolated hearts included; ventricular premature beats (VPBs), bigeminy, trigemini, potentiated contraction and ventricular tachycardia (VT) to form a five-point arrhythmia score classification which indicated severity of the arrhythmias (Supplementary Table 2). Three minutes of the trace recording for each drug treatment were analysed for abnormal ventricular contractions. Isolated hearts were also Langendorff-perfused without insertion of the balloon and treated with the same drug infusions or only KHB and immediately snap frozen in liquid nitrogen for analysis of protein *S*-nitrosylation.

**Table 1.** Echocrdiography parameters in WT and CaMKIIδ-C273S mice at 12 weeks.

| Parameter | WT | CAMKIIδ-C273S | Genotype effect (P value) |
| --- | --- | --- | --- |
| Animal weight (g) | 26.81 ± 1.11 | 30.06 ± 1.92 | 0.15 |
| HR (beats/min) | 419.9 ± 6.66 | 416.9 ± 11.34 | 0.81 |
| IVSd (mm) | 0.99 ± 0.02 | 1.00 ± 0.02 | 0.75 |
| IVSs (mm) | 1.29 ± 0.04 | 1.30 ± 0.03 | 0.90 |
| EF (%) | 79.48 ± 1.20 | 69.54 ± 2.65 | 0.001** |
| LVEDV (ml) | 0.07 ± 0.02 | 0.11 ± 0.01 | <0.0001**** |
| LVESV (ml) | 0.01 ± 0.00 | 0.03 ± 0.00 | <0.0001**** |
| R-R Interval (ms) | 144.2 ± 2.33 | 146.1 ± 4.05 | 0.67 |
| E/A Ratio | 2.31 ± 0.06 | 1.93 ± 0.07 | 0.0003*** |

HR, heart rate; IVSD, interventricular septum diastolic thickness; IVSS, interventricular septum systolic thickness; FS, Fractional shortening; EF, Ejection Fraction; LVEDV, left ventricular end diastolic volume; LVESV, left ventricular end systolic volume. Data are expressed as mean ± SEM. Independent t-test was used to compare between WT (n=12) and C273S (n=11) values, \* P < 0.05, \*\* P < 0.01, \*\*\* P < 0.001, \*\*\*\* P < 0.0001, WT vs C273S

### Experimental solutions

All drugs were prepared fresh daily or from frozen aliquots. ISO (100 nM, Merck, Germany) was used to initiate the β-AR signalling pathway. The NO donor GSNO (150 μM, Cayman Chemical, USA) was used to induce *S*-nitrosylation in cardiomyocytes ^19^. Sodium nitroprusside (SNP; 200 μM, Sigma, USA) was also tested as a clinical NO donor. The soluble guanylyl cyclase (sGC) inhibitor ODQ (10μM, Tocris, UK) was used to prevent activation of the NO-sGC-CGMP pathway.

### Modified Biotin switch for *S*-nitrosylated protein detection

Frozen whole heart samples were homogenized at 4°C with a Precellys Evolution homogenizer (Bertin) in homogenization buffer containing (in mM); 300 sucrose, 250 HEPES-NaOH, pH 7.7, 1 EDTA, 100 neocuproine, 0.5% Triton-X 100 and 25 N-ethylmaleimide (NEM, Sigma). One tablet of EDTA-free protease inhibitor (Roche) was added just before use. All buffers were freshly prepared before use and subsequent procedures were carried out in the dark to prevent degradation of light-sensitive *S*-nitrosylated proteins. Homogenates were incubated on ice for 15 min and centrifuged at 14000 rcf for 10 min at 4°C. The supernatant was recovered as total crude homogenate, aliquoted and stored at −80°C. Total protein concentration was determined using the Bradford protein assay.

Homogenates (100 μg protein) were diluted in HEN buffer containing (in mM): 250 HEPES-NaOH, pH 7.7, 1 EDTA, 0.1 neocouproine, 2% SDS, 20 NEM and incubated and oscillated at 80 rpm, 50°C for 40 min. Excess NEM was removed via cold acetone precipitation (−20°C for 20 min) and the pellet was collected via centrifugation at 10,000 rcf for 10 min at 4°C. Between each precipitation, the sample pellet was air-dried to ensure total removal of acetone. Samples were then resuspended in HEN with 1% SDS (wt./vol) and incubated with ascorbate (Sigma) and 4 μM DyLight Maleimide 800 (Thermo Fisher) to reduce *S*-nitrosylated cysteine residues and label with the maleimide group. Excess dye was removed via cold acetone precipitation (−20°C for 20 min) and samples were resuspended in 50μL of 1X reducing buffer (Invitrogen loading dye and 5% β-mercaptoethanol), heated for 5 mins (95°C) and separated on 4-12% SDS PAGE gel at 75 V for 20 min and 150 V for 80 min. The gel was then rinsed with deionized water and fluorescence was visualized at 800 nm using an iBright Imaging System (Thermo Fisher).

### Electrocardiogram recordings

Electrocardiograms (ECGs) were recorded from mice under 1.5-2 % isoflurane delivered via a nose cone with 100% O_2_ at 0.5 L/min. Negative, positive and earth electrodes were inserted subcutaneously with acupuncture needles into the right front limb, left back limb and right back limb respectively. ECG measurements were acquired from lead II connected to a Powerlab and recorded in LabChart 8.1(ADInstruments). Recordings were made for 10 minutes with the last 5 minutes used for analysis performed in LabChart.

### Data Analysis

All individual data points are shown for cardiomyocyte Ca^2+^ and isolated heart parameters along with the mean ± SEM. For results reported in the text, the data are mean ± SD. Isolated heart perfusion data were analysed on Lab Chart 8.1 (ADInstruments). Statistical analysis was performed using Prism 9 (GraphPad). ANOVA with a Sidak’s multiple comparisons test (two conditions) or Tukey’s multiple comparisons test was used for analysis where more than two conditions were tested on any given cardiomyocyte. Post hoc analysis with Fisher’s least significant difference (LSD) test was also done for the groups of Langendorff-perfused hearts. Differences in the fraction of cardiomyocytes displaying Ca^2+^ waves were determined by a Chi-squared test (between groups) or a McNemar’s test (paired data). Western blot, fibrosis and cell dimension and ECG data were compared using unpaired t-tests. Significant differences are shown for P≤0.05 (*), P<0.01 (**), P<0.001(***) and P<0.0001(****).

## Results

### NO donor does not alter baseline Ca^2+^ transient properties

To induce *S*-nitrosylation of target proteins we used the NO donor GSNO, which spontaneously releases NO into the perfusate (Figure 1A). [NO] peaked in KRH buffer at 0.54 ± 0.06 μM after 10 min and remained stable over the duration of our experimental timeframe (Figure 1B). Ventricular cardiomyocytes were isolated from WT and CaMKIIδ-KO mice which had undetectable expression of CaMKIIδ (Figure 1C). The cardiomyocytes were exposed to 150 μM GSNO according to the protocol outlined in Figure 1D. There were no observed effects of GSNO on Ca^2+^ transient amplitude (Figure 1E) or time constant Tau of Ca^2+^ decay (Figure 1F) in either WT or CaMKIIδ-KO myocytes. We also measured Ca^2+^ sparks in unpaced cardiomyocytes with confocal linescan imaging (Figure 1G). We observed no effect of GSNO on Ca^2+^ spark frequency in cardiomyocytes of either genotype (Figure 1H). There was also no effect of GSNO on SR Ca^2+^ content (Supplementary Figure 1A). Taken together, our data show that at the baseline low frequency stimulation used here, there is no major difference in electrically evoked or spontaneous Ca^2+^ transients in WT vs. CaMKIIδ-KO, nor does exposure to GSNO alone alter these properties.

**Figure 1.**
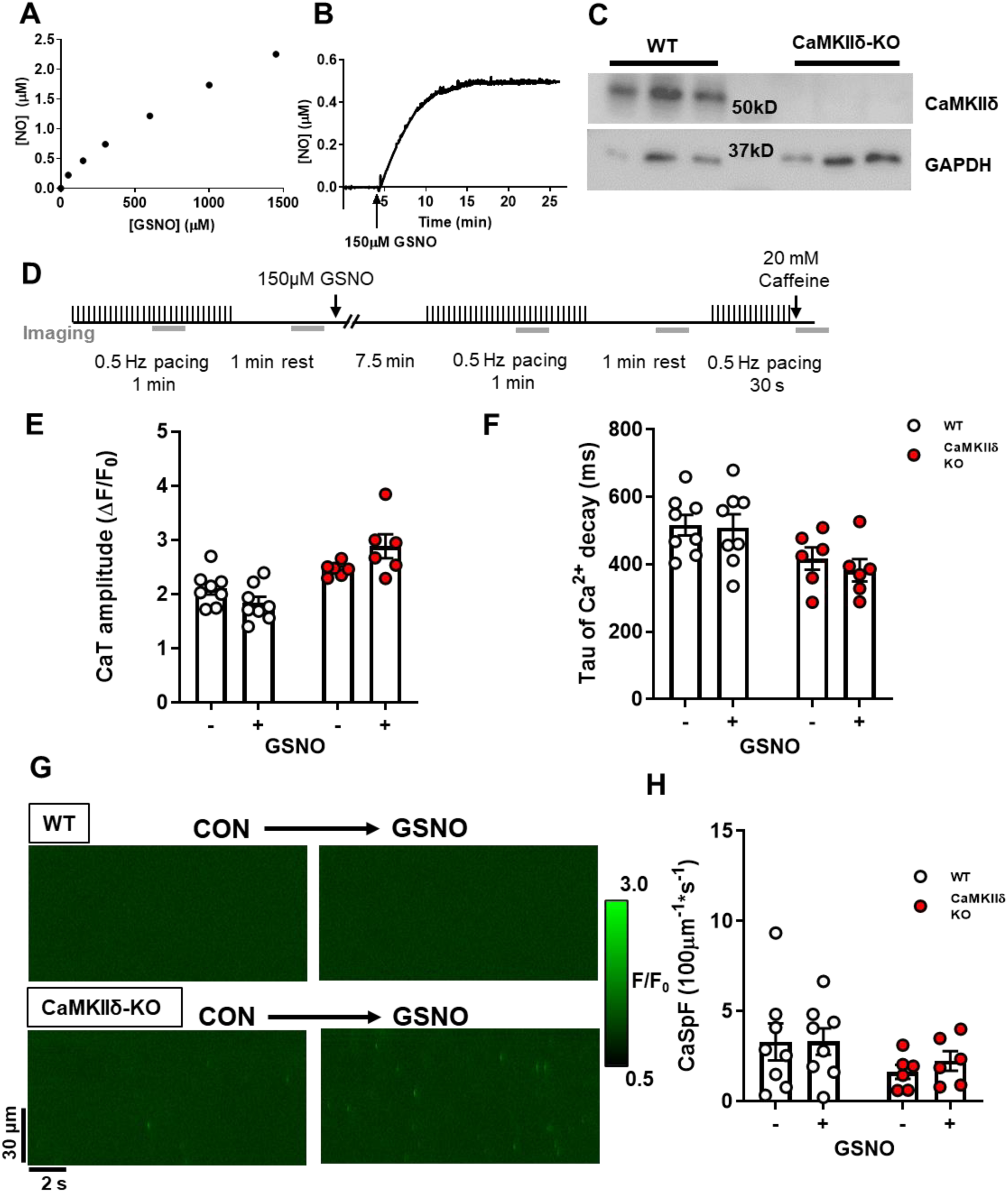
Cardiomyocyte Ca^2+^ transients and sparks with nitric oxide donor GSNO. (**A**) NO liberated from increasing GSNO concentrations in buffer was measured using a NO electrode. (**B**) The kinetics of NO release following the addition of 150 μM GSNO to buffer. **(C)** Western blot of ventricle tissue samples from WT and CaMKIIδ KO mice (n=3 hearts) showing loss of CaMKIIδ protein expression in CaMKIIδ-KO vs. WT hearts (with GAPDH loading controls). (**D**) WT or CaMKIIδ-KO cardiomyocytes were treated with 150 μM GSNO and paced at 0.5 Hz. There was no change in Ca^2+^ transient amplitude (**E**) or decay kinetics (**F**). In quiescent cardiomyocytes, Ca^2+^ sparks were measured from line-scan images (**G**) and there was also no effect of GSNO on Ca^2+^ spark frequency in either WT or CaMKIIδ-KO cardiomyocytes (**H**), (WT n=8 cells, N=3 hearts; CaMKIIδ-KO n=8 cells, N=3 hearts).

### CaMKIIδ-KO cardiomyocytes produce fewer Ca^2+^ sparks during β-adrenergic stimulation

Next we increased myocyte Ca^2+^ transients with exposure to 100 nM ISO, which is expected to strongly promote CaMKIIδ activation (Figure 2A). Representative confocal line-scans from Fluo-4-AM loaded cardiomyocytes isolated from WT and CaMKIIδ-KO hearts are shown in Figure 2B. Fluo-4 fluorescence was normalized to baseline (F_0_) to determine the Ca^2+^ transient characteristics in WT and CaMKIIδ-KO cardiomyocytes during ISO exposure (Figure 2C). ISO induced a three-fold increase in Ca^2+^ transient amplitude (Figure 2D), while Ca^2+^ transeint decay was twice as fast as in control conditions in both WT and CaMKIIδ-KO cardiomyocytes (Figure 2E). The SR Ca^2+^ content, as measured by peak Ca^2+^ release after caffeine application, was increased by ISO and did not differ between mouse genotypes (Supplementary Figure 1B; p = 0.03 for overall ISO effect). ISO increased Ca^2+^ spark frequency (WT: p < 0.0001; CaMKIIδ-KO: p = 0.025) and amplitude (WT control 0.66 ± 0.49 ΔF/F_0_ vs ISO 1.21 ± 0.61 ΔF/F_0_ p < 0.0001; CaMKIIδ-KO control 0.58 ± 0.15 ΔF/F_0_ vs ISO 0.83 ± 0.29 ΔF/F_0_: p = 0.019) in both WT and CaMKIIδ-KO cardiomyocytes; however, the increase in spark frequency was significantly attenuated in the CaMKIIδ-KO cardiomyocytes (p<0.0001; Figure 2F-G) and there was a trend towards a smaller Ca^2+^ spark amplitude (p = 0.057). Thus, while CaMKIIδ deletion did not prevent the effect of ISO on overall Ca^2+^ transient properties, it limited spontaneous Ca^2+^ release during β-AR stimulation.

**Figure 2.**
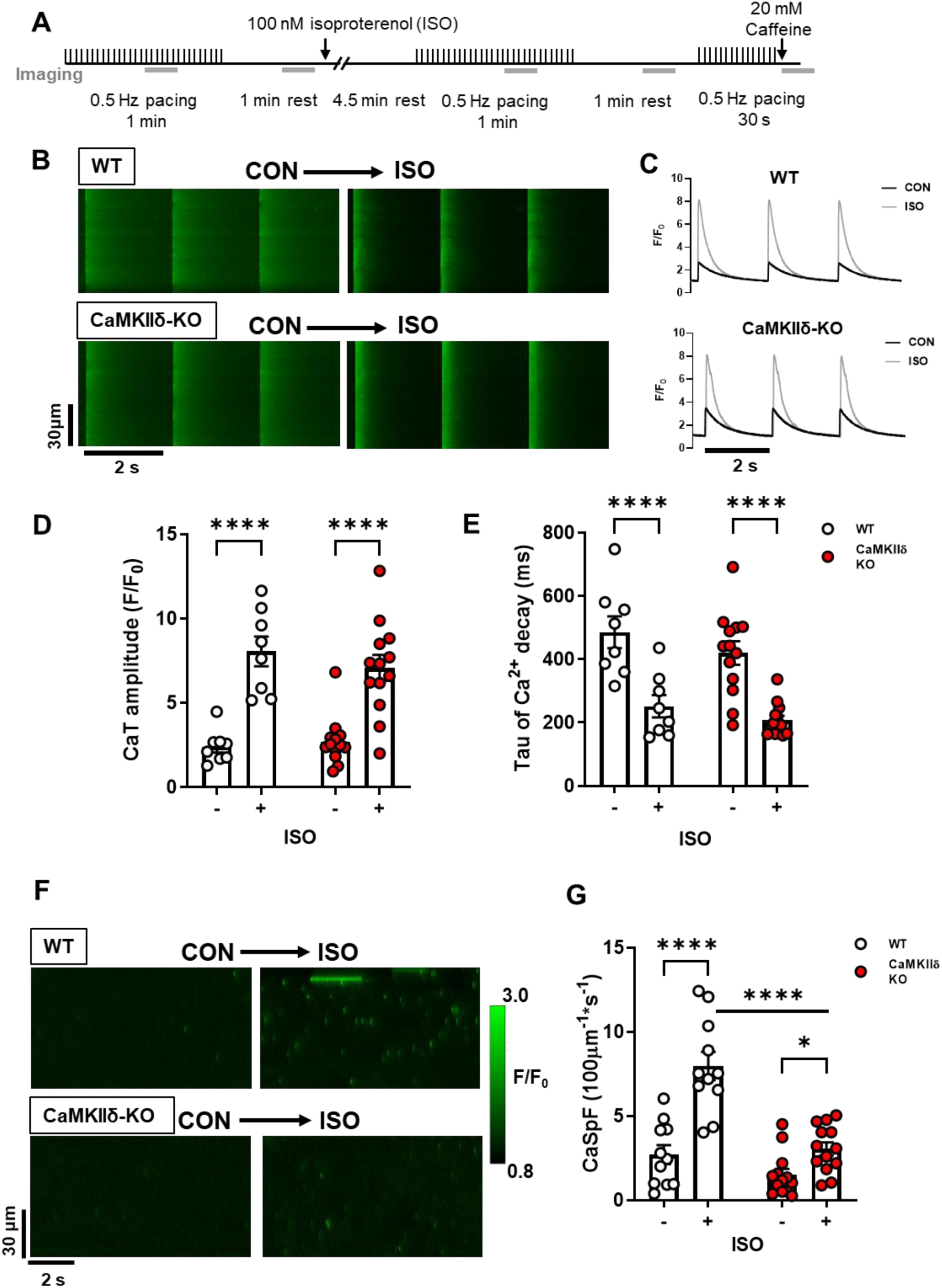
Cardiomyocyte Ca^2+^ transient and spark properties in response to β-adrenergic agonist isoproterenol. (**A**) Experimental protocol used for Ca^2+^ imaging, the grey bars represent when line-scan images were acquired. Representative line-scans during pacing (**B**) and resultant Ca^2+^ transients (**C**) from a WT and CaMKIIδ-KO cardiomyocyte stimulated at 0.5 Hz under control conditions and with 100 nM ISO. ISO increased Ca^2+^ transient amplitude (**D**) and accelerated decay (**E**) in both WT and CaMKIIδ-KO cardiomyocytes to a similar degree (WT n=8 cells, N=2 hearts; CaMKIIδ-KO n=13 cells, N=4 hearts). (**F**) Representative line-scans from a WT and CaMKIIδ-KO cardiomyocyte show an increase in the number of Ca^2+^ sparks following exposure to ISO. Mean data show the ISO-induced increase in Ca^2+^ spark frequency (CaSpF) was attenuated in the CaMKIIδ-KO cardiomyocytes (**G**) (WT n=11 cells, N=2 hearts; CaMKIIδ-KO n=13 cells, N=4 heats).

### The Cys-273 site on CaMKIIδ attenuates SR Ca^2+^ release in response to NO and ISO

We previously showed that *S*-nitrosylation of the Cys-273 site of CaMKIIδ *in vitro* can prevent activation of the kinase in response to increased Ca^2+^/CaM ^19^. We hypothesized that this mechanism might protect myocytes from the development of arrhythmogenic events by supressing CaMKIIδ activity during transient periods of nitrosative stress. To test this hypothesis, we generated knock-in mice that lack the inhibitory *S*-nitrosylation site (CaMKIIδ-C273S; Figure 3A-C). The CaMKIIδ-C273S mice had normal CaMKIIδ and RyR2 expression however, there was an increased expression of SERCA2A (Figure 3D-H) in the ventricle. There was no evidence of ventricular fibrosis (Figure 3G &I) or cardiomyocyte hypertrophy (Figure 3 J-L) in the CaMKIIδ-C273S mouse hearts compared to WT. Isolated ventricular myocytes from WT and CaMKIIδ-C273S mice were treated with 150 μM GSNO for 7 min immediately prior to wash-in of 100 nM ISO without GSNO. Pre-incubation of cardiomyocytes with GSNO had no effect on baseline Ca^2+^ transient amplitude (Figure 4A; WT: p > 0.999; CaMKIIδ-C273S: p = 0.838) or time constant of [Ca^2+^]_i_ decay (Figure 4B; WT: p > 0.9999; CaMKIIδ-C273S: p = 0.301) relative to cardiomyocytes in normal control KRH buffer. Moreover, WT cardiomyocytes pre-treated with GSNO showed a typical Ca^2+^ transient response to ISO, with Ca^2+^ transient amplitude (p > 0.999) and decay (p > 0.999) similar in magnitude to control WT cardiomyocytes not subject to GSNO pre-treatment (Figure 4A-B). Notably, cardiomyocytes from CaMKIIδ-C273S mice pre-treated with GSNO had larger ISO induced Ca^2+^ transients compared to WT cardiomyocytes (Figure 4A, bar 4 vs. bar 8; p < 0.0001) and in comparison, to CaMKIIδ-C273S cardiomyocytes not pre-treated with GSNO (Figure 4A, bar 6 vs. bar 8; p = 0.0002). These data indicate that genetic ablation of the inhibitory C273 *S*-nitrosylation site increases the effects of CaMKIIδ on SR Ca^2+^ release in the presence of NO and is consistent with our hypothesis that *S*-nitrosylation of the C273 site is protective in limiting CaMKIIδ activation and its effects on RyR2-related Ca^2+^ transient properties.

**Figure 3.**
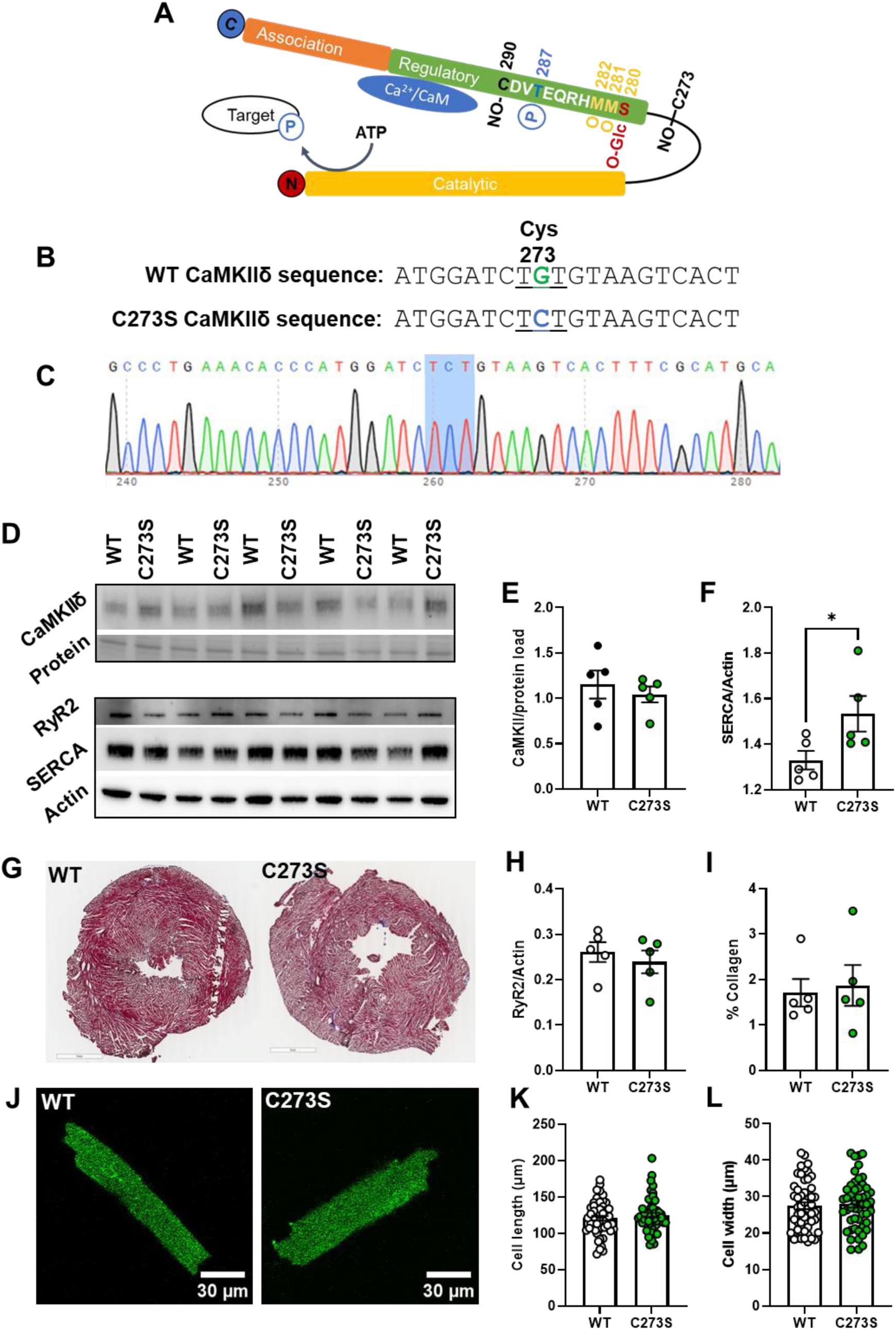
Characterization of the CaMKIIδ-C273S knock-in mouse model. Schematic of the CaMKIIδ monomer with the residue positions of various post-translational modifications within the regulatory domain including S-nitrosylation (NO-), phosphorylation (P), oxidation (O) and O-GlcNAc modification which are known to alter CaMKII activity (**A**). A mouse model was generated using CRISPR/cas9 causing a single point mutation in the cystine codon at position 273, resulting in replacement with a serine residue which cannot be S-nitrosylated (**B**). Example of a CaMKIIδ-C273S offspring confirmed with genotyping of ear notch (**C**). Protein expression in ventricular tissue (N = 5 hearts) was measured using western blots (**D**) for CaMKIIδ (**E**), SERCA (**F**) RyR (**H**), and normalized to actin or protein load. Ventricular fibrosis was measured in fixed and stained tissue (**G**) by quantifying collagen content (**I**). Representative cardiomyocytes are shown in J for measurement of cell length (**K**) and width (**L**), WT: n = 52, N = 7 hearts; CaMKIIδ-C273S: n=50, N=8 hearts.

**Figure 4.**
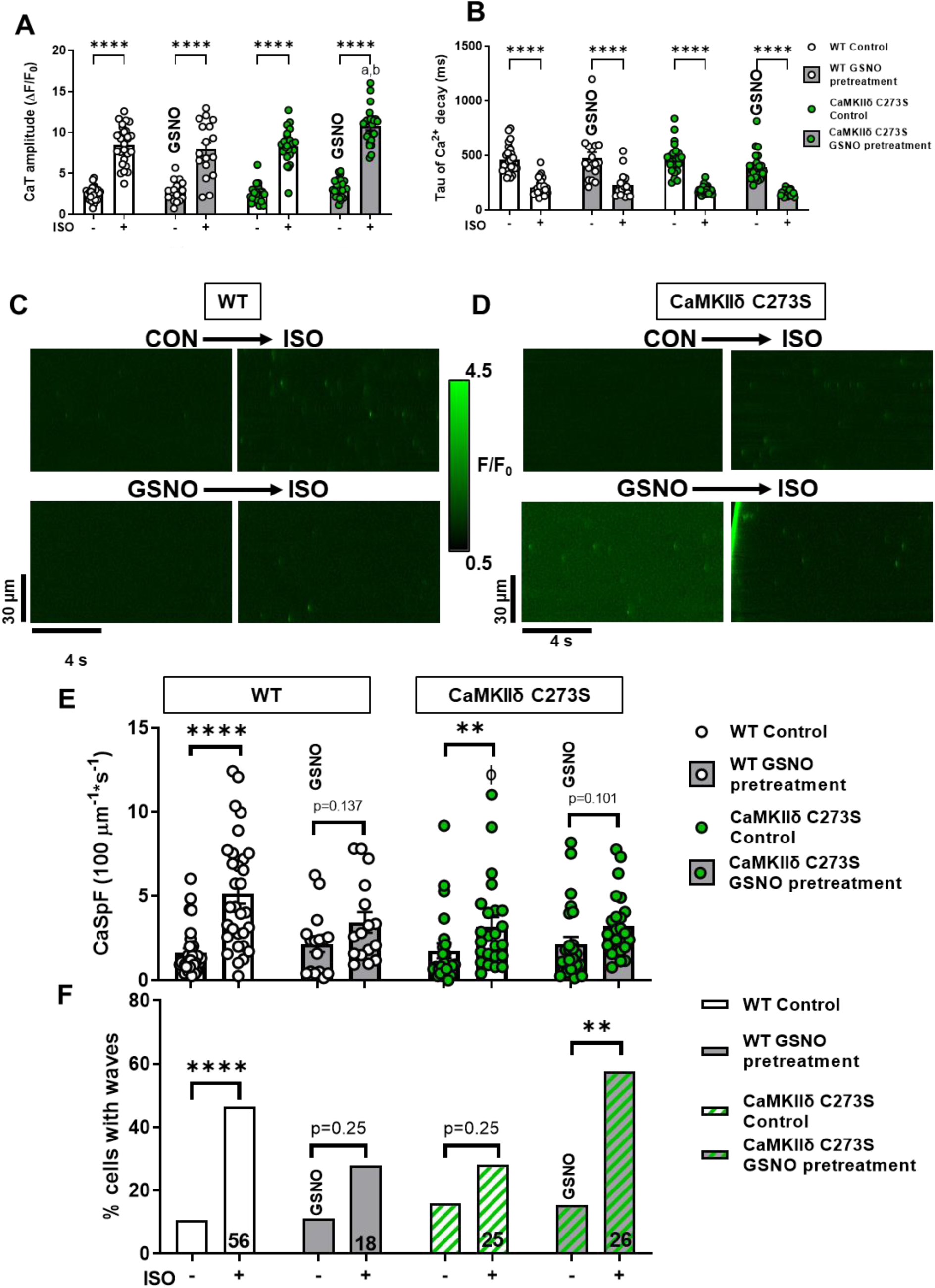
Pre-treatment with GSNO enhances Ca^2+^ transient response to isoproterenol in CaMKIIδ-C273S cardiomyocytes. Ca^2+^ transients from WT and CaMKIIδ C273S cardiomyocytes from were recorded at 0.5 Hz under control conditions and with 100 nM ISO or with pre-treatment of 150 μM GSNO prior to ISO exposure. Mean Ca^2+^ transient amplitude (**A**) and decay (**B**) data in response to ISO with control KRH buffer (white bars; WT: n=28 cells, N=14 hearts; CaMKIIδ-C273S: n=24 cells, N=14 hearts) or when pre-treated with GSNO (grey bars; WT: n=18 cells, N=7 hearts; CaMKIIδ-C273S: n=24, N=9 hearts). Representative line-scans from quiescent WT (**C**) and CaMKIIδ-C273S (**D**) cardiomyocytes under control conditions and with ISO or with pre-treatment of GSNO prior to ISO exposure. Mean Ca^2+^ spark frequency (**E**) data are shown for cardiomyocytes in response to ISO with control buffer (white bars; WT: n=31 cells, N=14 hearts; CaMKIIδ-C273S: n=24 cells, N=9 hearts) or when pre-treated with GSNO (grey bars; WT: n=16 cells, N=7 hearts; CaMKIIδ-C273S: n=25 cells, N=9 hearts). Percentage of cardiomyocytes exhibiting Ca^2+^ waves in quiescent WT and CaMKIIδ-C273S cardiomyocytes (**F**). ɸ p<0.05 compared to WT ISO, ^a^p = 0.0002 vs CaMKIIδ-C273S ISO, ^b^p <0.0001 vs WT ISO with GSNO pre-treatment.

Representative line-scans to detect spontaneous Ca^2+^ leak with GSNO pre-treatment followed by ISO are shown for WT (Figure 4C) and CaMKIIδ-C273S (Figure 4D) cardiomyocytes. Pre-treatment with GSNO did not alter baseline Ca^2+^ spark frequency for either WT (p = 0.9996) or CaMKIIδ-C273S (p >0.999) cardiomyocytes (Figure 4E). ISO increased the frequency of Ca^2+^ sparks in both WT and CaMKIIδ-C273S cardiomyocytes. However, this increase was suppressed in WT cardiomyocytes pre-treated with GSNO (Figure 4E, bar 3 to 4 vs. 1 to 2). Surprisingly, in the C273S myocytes we also observed a suppression of ISO-induced Ca^2+^ sparks (Figure 4E, bar 7 to 8 vs. 5 to 6) where the inhibitory effect of *S*-nitrosylation at Cys-273 was expected to be prevented.

To further interrogate potential arrhythmogenic signalling, we quantified larger propagating pro-arrhythmic Ca^2+^ release events known as Ca^2+^ waves. ISO induced an increase in Ca^2+^ waves in WT cardiomyocytes, but in WT this effect was supressed by GSNO pre-treatment (Figure 4F, left). In contrast, CaMKIIδ-C273S cardiomyocytes pre-treated with GSNO robustly increased ISO-induced Ca^2+^ waves (Figure 4F, right), suggesting that the inhibitory Cys-273 *S*-nitrosylation site is important for preventing Ca^2+^ waves that can trigger action potentials and arrhythmias. Notably, the occurrence of Ca^2+^ waves causes relative unloading of the SR (Supplemental Fig 1C), which would indirectly limit Ca^2+^ spark frequency (Figure 4E bar 7 vs. 8). We repeated the experiments with SNP, a NO donor used in clinical settings, which demonstrated a similar attenuation of Ca^2+^ sparks in the WT cardiomyocytes (as for GSNO pre-treatment) which was lost in the CaMKIIδ-C273S cardiomyocytes, without altering the Ca^2+^ transient response to ISO (Figure 5A-D). Preventing activation of the NO-sGC-cGMP pathway with sGC inhibitor ODQ also did not alter the response of the WT cardiomyocytes, particularly the Ca^2+^ spark effects of ISO (Figure 5E-H). We infer that the SNP (and GSNO) mediated protection against ISO-induced arrhythmogenic SR Ca release events is mediated mainly by *S*-nitrosylation of CaMKIIδ-C273.

**Figure 5.**
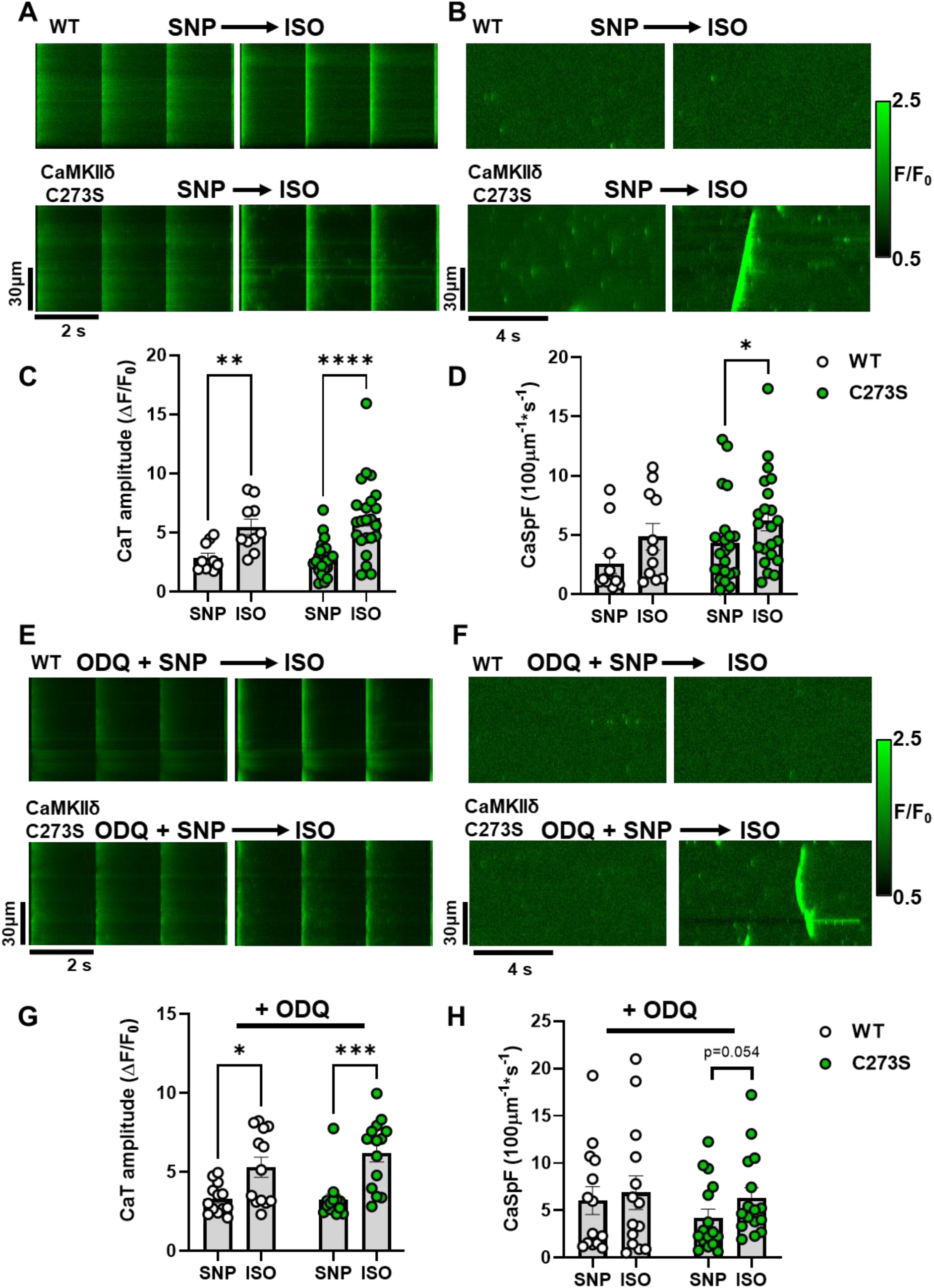
Pre-treatment with clinical NO donor SNP and sGC inhibitor elicits similar protection against isoproterenol induced Ca^2+^ sparks. Ca^2+^ transients (**A**) and sparks (**B**) from WT and CaMKIIδ-C273S cardiomyocyte were recorded at the end of a 7-minute incubation with SNP (200 μM) followed by wash-in of 100 nM ISO. Mean Ca^2+^ transient amplitude and spark frequency data are shown in C and D respectively (WT: n=10 cells, N=3 hearts; CaMKIIδ-C273S: n=23 cells, N=6 hearts). The same experiments were performed following a pre-treatment with sGC inhibitor ODQ (10 μM; 20 min incubation) with representative line-scans (**E**-**F**) and mean data (**G**-**H**) plotted for WT (n=13 cells, N=3 hearts) and CaMKIIδ-C273S (n=14 cells, N=4 hearts) cardiomyocytes.

### NO treatment after β-AR activation does not alter Ca^2+^ transients or sparks

Previous literature has demonstrated a role for autonomous activation of CaMKIIδ by *S*-nitrosylation at Cys-290 ^19^ and enhanced Ca^2+^ release events mediated by CaMKII ^13^. Due to the position of the Cys-290 in the CaMKIIδ regulatory domain adjacent to the Thr-287 autophosphorylation site, the *S*-nitrosylation site is unlikely to be available under basal conditions when the kinase is auto-inhibited by regulatory domain binding to the catalytic domain ^19^. Therefore, we tested the effect of GSNO following ISO treatment when the CaMKIIδ activation state is increased in cardiomyocytes ^26^. We also examined the effects of ISO and GSNO on ventricular myocytes isolated from CaMKIIδ-C290A knock-in mice, in addition to WT and CaMKIIδ-C273S animals.

Cardiomyocytes underwent wash-in of 100 nM ISO for 5 min followed by washout with control buffer, or buffer containing 150 μM GSNO. Ca^2+^ transients were measured at baseline, during ISO wash-in, and following washout with GSNO buffer (Figure 6A). ISO increased Ca^2+^ transient amplitude and accelerated decay in WT and CaMKIIδ-C273S cardiomyocytes as expected, and these effects were sustained during washout in the presence of GSNO (Figure 6B-C, white and green symbols). Similar Ca^2+^ transient results were observed in CaMKIIδ-C290A cardiomyocytes, both before and after treatment with GSNO (Figure 6C, blue symbols).

**Figure 6.**
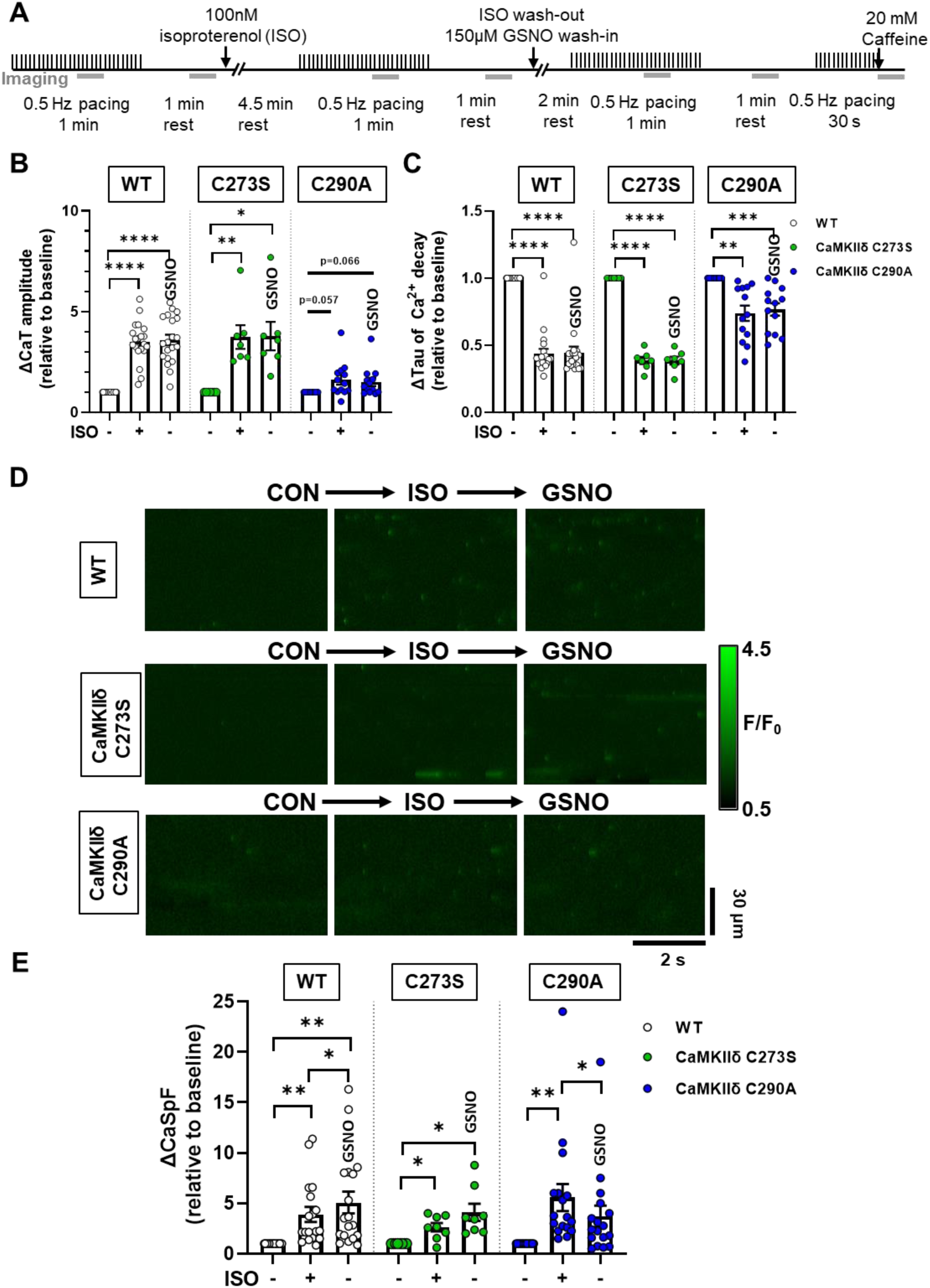
Effects of GSNO after ISO on Ca^2+^ transients and sparks in WT or CaMKIIδ-C273S and -C290A cardiomyocytes. WT, CaMKIIδ-C273S and -C290A myocytes were stimulated at 0.5 Hz and exposed to 100 nM ISO, then following ISO exposure myocytes were exposed to 150 μM GSNO. Experimental protocol used for Ca^2+^ imaging, the grey bars represent when line-scan images were acquired (**A**). Mean Ca^2+^ transient amplitude (**B**) and decay (**C**) data are shown for cardiomyocytes in response to ISO and wash-out with GSNO (WT: n=20 cells, N=9 hearts; CaMKIIδ C273S: n=7 cells, N=5 hearts; CaMKIIδ-C290A: n=13 cells, N=5 hearts). Representative line-scans from quiescent WT, CaMKIIδ-C273S and -C290A cardiomyocytes under indicated conditions (**D**). Mean Ca^2+^ spark frequency data (**E**) are shown for cardiomyocytes in response to ISO with washout with GSNO (WT: n=18 cells, N=9 hearts; CaMKIIδ C273S: n=8 cells, N=5 hearts; CaMKIIδ-C290A: n=17 cells, N=5 hearts). All data are normalized to baseline control before addition of ISO.

When spontaneous Ca^2+^ release was measured in quiescent cardiomyocytes (Figure 6D), ISO resulted in an increase in the Ca^2+^ spark frequency (Figure 6E) for all three genotypes. Addition of GSNO during the ISO washout prevented Ca^2+^ spark frequency from returning to baseline in both the WT and CaMKIIδ-C273S myocytes. However, myocytes from the CaMKIIδ-C290A mice had significantly reduced Ca^2+^ spark frequency during the ISO washout, despite the presence of GSNO. These data are consistent with the hypothesis that ISO-induced CaMKIIδ activation regulates Ca^2+^ handling in myocytes, and subsequent exposure to GSNO causes *S-*nitrosylation at the Cys-290 site, thereby prolonging the enhancement of CaMKIIδ activity and Ca^2+^ sparks.

### Knock-in CaMKIIδ-C273S mice have impaired cardiac function and altered ECG characteristics at 12 weeks of age

The myocyte Ca^2+^ spark and wave data (Figure 4) suggest that the CaMKIIδ-C273 *S*-nitrosylation site might offer basal protection from stress associated with sympathetic activation of β-adrenergic receptors (i.e. that is lost in CaMKIIδ-C273S mice). To test whether there is chronic cardiac adaptation in terms of cardiac function, we measured these parameters in anesthetized WT and CaMKIIδ-C273S mice at 12 weeks of age (Table 1). The echocardiography data indicated unaltered body weight, heart rate, and septal thickness between the groups. However, both end diastolic and systolic volumes of the LV were significantly increased in the CaMKIIδ-C273S mice, with reduced fractional shortening and ejection fraction. The ratio of early to late ventricular filling velocities (E/A ratio) was also significantly impaired in the CaMKIIδ-C273S hearts. Taken together, these data show that the lack of one *S*-nitrosylation site in the CaMKIIδ-C273S mice results in a modest reduction in both systolic and diastolic function, even absent a significant cardiac challenge.

We also measured conduction characteristics through the hearts of anesthetized WT (Figure 7A) and CaMKIIδ-C273S (Figure 7B) using ECG. Even at 12 weeks of age, we observed significant differences in a number of ECG parameters (Figure 7C-K) for the CaMKIIδ-C273S mice, including prolongation of P wave duration and P-R interval, as well as increased Q, R, and S wave amplitudes. There was also a trend towards increased T wave amplitude in the CaMKIIδ-C273S mice (p=0.058). In addition to these baseline alterations to conduction, we observed spontaneous arrhythmic events in the ECG traces of the CaMKIIδ-C273S mice. Figure 7L shows an example of periodic spontaneous variability in heart rate, which was observed in several of the CaMKIIδ-C273S mice but none of the WT mice (Figure 7M), leading to occasional periods of bradycardia (Figure 7N). These arrhythmic events were excluded from our analysis of ECG parameters (Figure 7C-K), as this would have greatly exaggerated the differences in these values. Nonetheless, our observations that conduction is altered and spontaneous arrhythmic events are occurring in the CaMKIIδ-C273S mice suggest that the Cys-273 site on CaMKIIδ is playing a critical protective role in the heart.

**Figure 7.**
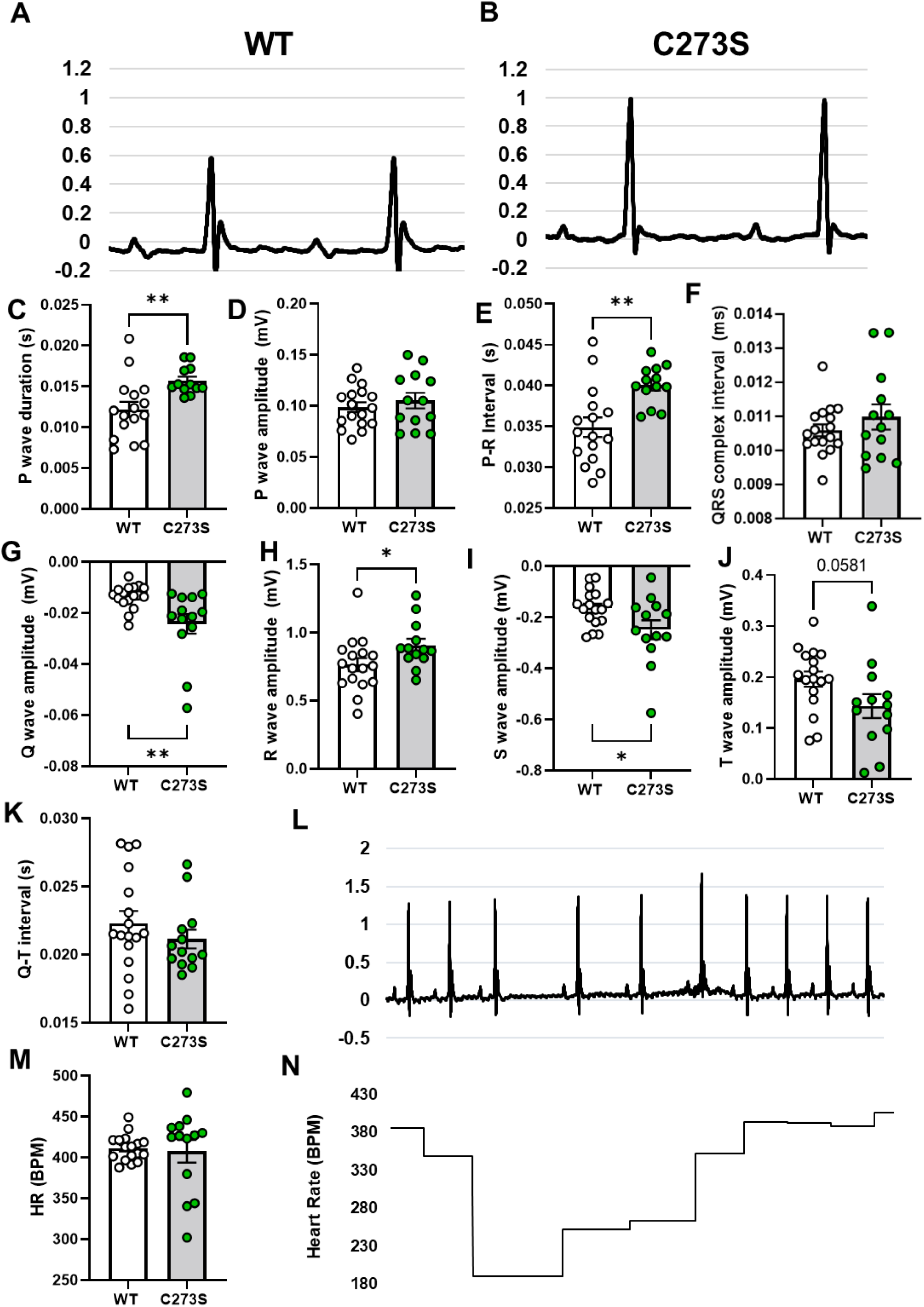
Electrocardiogram characteristics for WT and CaMKIIδ-C273S mice. Representative electrocardiograms from a WT (**A**) and CaMKIIδ-C273S (**B**) mouse under anaesthetic and mean data for waveform amplitudes and durations (**C-K**). N = 13-17 per group. ECG traces for some of the CaMKIIδ-C273S mice showed spontaneous arrhythmic events (**L**). These events led to variability in total heart rate (**M**) and acute periods of bradycardia (**N**).

### NO mediates arrhythmogenic response to β-AR stress in Langendorff-perfused mouse hearts

Our observation that GSNO pre-treatment could suppress ISO-induced Ca^2+^ sparks in isolated cardiomyocytes, motivated us to test the arrhythmogenic consequences at the whole heart level using Langendorff-perfused mouse hearts (Figure 8A). To ensure that the GSNO treatment was sufficient to induce *S*-nitrosylation during Langendorff-perfusion, we snap-froze hearts after perfusion and used a modified biotin switch assay to measure total *S*-nitrosylation (Figure 8B). Hearts that received GSNO after ISO had a significant increase in total *S*-nitrosylation (Figure 8C, red vs. white bar; p<0.01), and this effect was partially reversed when GSNO was followed by 10 min ISO perfusion (Figure 8C, white vs. blue bar; p = 0.12).

**Figure 8.**
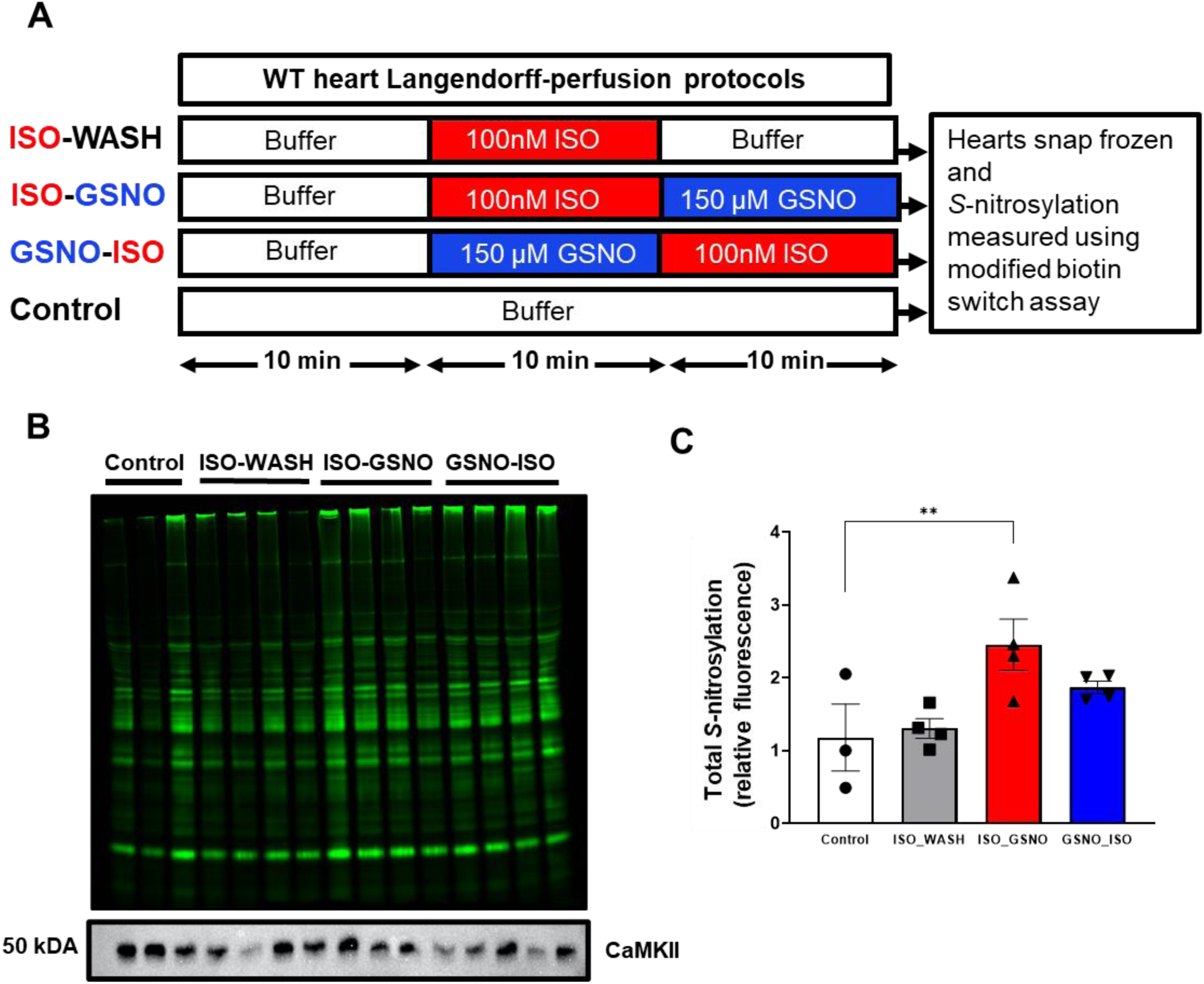
Whole mouse hearts exposed to ISO and GSNO have increased *S*-nitrosylation of cardiac proteins. (**A**) WT mouse hearts undergoing Langendorff-perfusion were treated with ISO and GSNO in varying order, snap-frozen and lysed, followed by a modified biotin switch assay to measure total protein *S*-nitrosylation. (**B**) Total cardiac protein *S*-nitrosylation in each lysate. (**C**) Treatment with ISO followed by GSNO (red bar) resulted in a significant increase in total cardiac protein *S*-nitrosylation compared to control (white bar, n=3-4 hearts per group).

We observed different types of arrhythmias in the isolated hearts treated with ISO and GSNO (Figure 9A). Treatment with 100 nM ISO induced a significant increase in total arrhythmic events (Figure 9B) and arrhythmia score (Figure 9C) in both the WT and CaMKIIδ-C273S hearts. After ISO washout, there was no significant difference in arrhythmic events compared to baseline for the WT or CaMKIIδ-C273S hearts. However, the arrhythmia score remained elevated above baseline for both groups even after ISO washout, indicating that some ISO-induced sensitization of the hearts persisted 10 min after washout. We then repeated the experiment with GSNO added to the perfusate during the ISO washout (analogous to Figure 6) to test whether GSNO would stabilize the arrhythmic phenotype after the ISO stress. GSNO treatment did prolong the increase in arrhythmias during the ISO washout phase for both WT and CaMKIIδ-C273S hearts (Figure 9D), and arrhythmia score remained significantly increased above baseline for both groups (Figure 9E). These data are consistent with our hypothesis that ISO-mediated stress activates CaMKIIδ to induce cardiac arrhythmia and that *S*-nitrosylation of activated CaMKIIδ (at Cys-290) can prolong this activation state, independent of Cys-273 availability.

**Figure 9.**
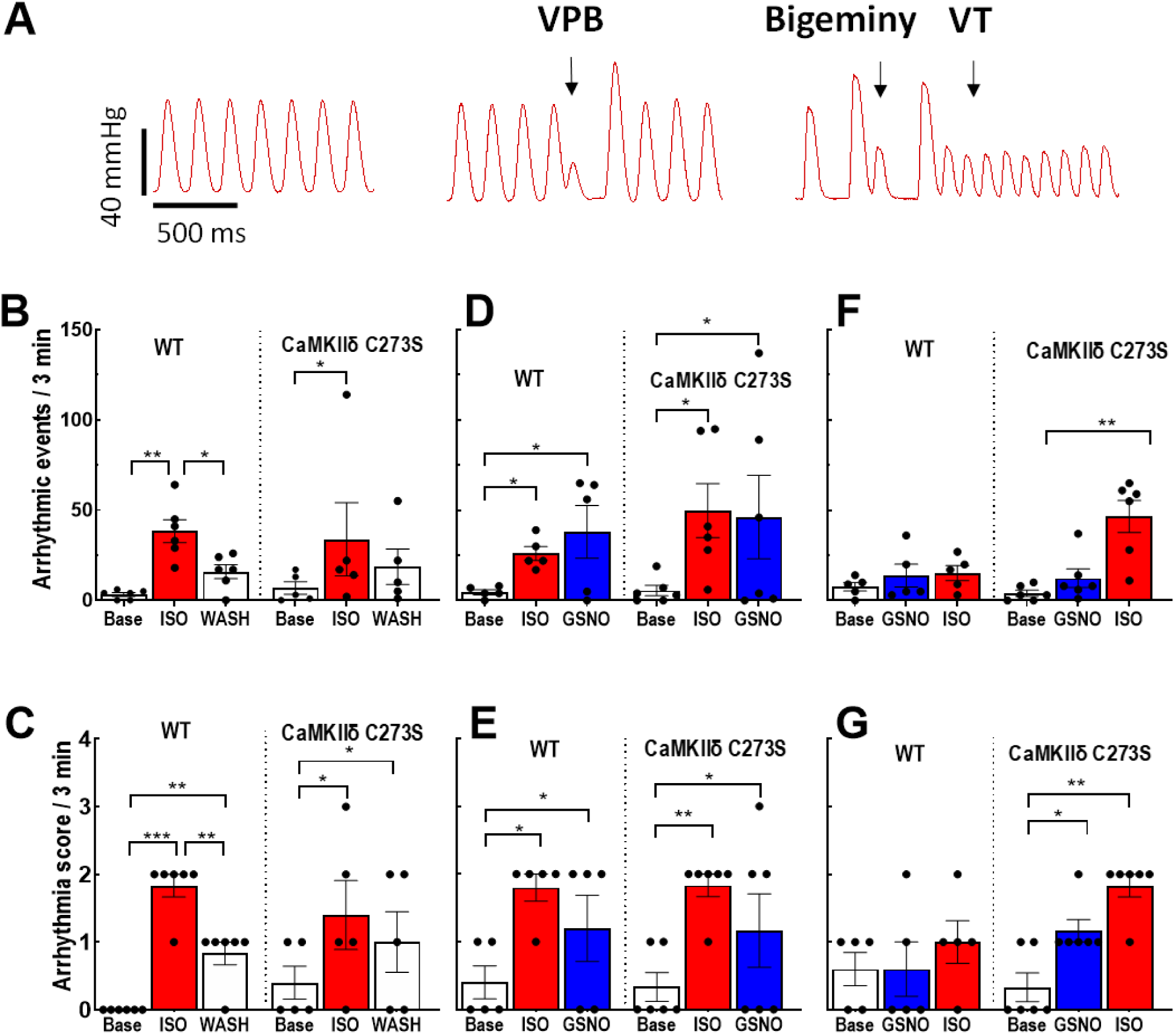
Pre-treatment with GSNO prevents ISO-induced arrhythmias in Langendorff-perfused hearts from WT, but not CaMKIIδ-C273S, mice. Arrhythmias were measured in Langendorff-perfused hearts following protocols outlined in Figure 7A. (**A)** Examples of arrhythmic events observed in the isolated hearts. We measured (**B**) quantity of arrhythmic events and (**C**) arrhythmia score during ISO perfusion only, (**D**) quantity of arrhythmic events and (**E**) arrhythmia score during ISO pre-treatment before GSNO, and (**F**) quantity of arrhythmic events and (**G**) arrhythmia score during GSNO pre-treatment before ISO. White bar = buffer, red bar = ISO, blue bar = GSNO. n = 5-6 hearts per group.

Our myocyte data showed that GSNO treatment prior to ISO exposure could limit arrhythmogenic Ca^2+^ wave activity in myocytes from WT but not CaMKIIδ-C273S mice (Figure 4F). To test this at the whole heart level, we exposed the hearts to GSNO first (10 min) before administration of ISO. Strikingly, WT mice were protected from ISO-induced arrhythmic events (or increases in arrhythmia score; Figure 9F-G), but this protection was not observed in the CaMKIIδ-C273S knock-in mice which lack the *S*-nitrosylation site that limits CaMKIIδ activation (Figure 9F). Moreover, in the CaMKIIδ-C273S mice, GSNO alone was sufficient to enhance the arrhythmia score (Figure 9G), a response that could be mediated through Cys-290 which promotes autonomous CaMKIIδ activation and would be unopposed in the absence of the inhibitory Cys-273 site. These data demonstrate that the Cys-273 site on CaMKIIδ is critical to the protective effect of GSNO pre-treatment with respect to ISO-induced arrhythmic response in the heart.

## Discussion

One main finding of this study was that NO donors promote CaMKIIδ-dependent spontaneous SR Ca^2+^ release following β-AR stimulation in cardiomyocytes and ventricular arrhythmias in the intact heart. Conversely, pre-treatment of cardiomyocytes or hearts with NO donors prior to ISO exposure limited the β-AR-induced increase in spontaneous Ca^2+^ release events and ventricular arrhythmias. Furthermore, this protective effect of NO pretreatment was mediated by the Cys-273 site on CaMKIIδ since it was attenuated when this site was mutated and not available for *S*-nitrosylation. Exposure to GSNO did not alter baseline Ca^2+^ handling or heart function. Notably, pre-treatment with GSNO conferred protection against β-AR stimulation of pathological Ca^2+^ leak and arrhythmias without compromising β-AR induced gain of function (enhanced Ca^2+^ transient amplitude and decay rate). These data provide a new understanding of how NO treatment influences β-AR signaling and cardiac function with both protective and pro-arrhythmic consequences depending on the timing of NO exposure.

While the β-AR signaling system is a vital arm of the ‘fight-or-flight’ response responsible for increasing cardiac output on demand, activation of the pathway can lead to aberrant Ca^2+^ release ^4,11,12,17^ and trigger arrhythmias ^27^. Activation of nitric oxide synthase (NOS) has been identified as a downstream signaling mechanism of β-AR stimulation, as endogenous NO levels increase following cardiomyocyte ISO exposure ^13,17^ and inhibition of NOS prevents an increase in Ca^2+^ leak with ISO ^11,13,17^. It has been proposed that NOS is activated following β-AR stimulation by a pathway independent of protein kinase A, apparently involving Epac and protein kinase B (Akt) ^11^. Here we found that ISO increased both Ca^2+^ spark frequency and amplitude in isolated cardiomyocytes, exposure of cardiomyocytes to GSNO alone under baseline conditions had no effect on triggered or untriggered Ca^2+^ events (Figures 2 and 4). This result was in contrast to the findings that GSNO (or SNP) increased Ca^2+^ sparks and arrhythmogenic waves in cardiomyocytes and was mediated by CaMKII nitrosylation and activation (rather than direct *S*-nitrosylation of RyR2) ^11,13^. As we’ve shown here, the directional effect of GSNO on CaMKII-dependent signaling is highly dependent on the basal state of CaMKII activation and whether Cys273 or Cys290 is the prime mediator, and that can depend on time, pacing rate, oxidative stress, Na^+^ and Ca^2+^ levels. That, and species differences ^28^ could readily explain baseline differences in basal GSNO effects. Indeed, cardiac CaMKIIδ activity modulates function of many myocyte targets ^18,29,30^, including ion channels associated with arrhythmias and excitation-contraction coupling such as RyR2 ^31^, SERCA (via phospholamban ^32^) and LTCC ^33^.

Under healthy resting conditions, CaMKIIδ is largely but not completely auto-inhibited and becomes progressively activated when intracellular Ca^2+^ levels rise and Ca^2+^/CaM is bound to the regulatory domain ^18^. Several post-translational modifications ^34–36^ of CaMKIIδ, including *S*-nitrosylation ^19^, enable an autonomously active form of the kinase that is associated with pathological Ca^2+^ mishandling and can be driven by high oxidative stress ^35^ or hyperglycaemia ^36^. The degree of CaMKIIδ *S*-nitrosylation is increased in cardiomyocytes treated with either GSNO ^13^ or ISO ^17^ as determined by immunoprecipitation of CaMKIIδ and probing with anti-*S*-nitrosylation antibodies. However, this method is not able to discriminate between the two Cys residues within the regulatory domain of CaMKIIδ that have opposite effects on CaMKIIδ activity ^19^. *S*-nitrosylation at Cys-290 causes autonomous activation of the kinase, consistent with the observation that NO exposure can enhance CaMKIIδ activity and increase Ca^2+^ sparks ^11,13^. Conversely, *S*-nitrosylation at Cys-273 inhibits CaMKIIδ by preventing Ca^2+^/CaM binding ^19^, suggesting a dual role for NO in mediating CaMKIIδ activity.

When cardiomyocytes are stimulated at 0.5 Hz, CaMKIIδ activation can be low ^26^, therefore, the Cys-290 residue within the regulatory region of CaMKIIδ may not be accessible for *S*-nitrosylation to induce autonomous activation ^19^. Stimulation of cardiomyocytes at a physiological rate (6-8 Hz in mice) would likely increase the sensitivity of the cells to the NO and CaMKIIδ mediated effects reported here in isolated cardiomyocytes; however, physiological rates are obtained in the isolated whole hearts and exposure to GSNO alone caused an increase in arrythmia score in CaMKIIδ-C273S hearts (Figure 9G). Under the baseline conditions for isolated cardiomyocytes used in this study, we propose that GSNO pre-treatment of cardiomyocytes results in *S*-nitrosylation of CaMKIIδ at Cys-273, which can prevent Ca^2+^/CaM dependent CaMKIIδ activation when subsequently stimulated with β-AR agonist ISO. We found that pre-treatment with GSNO prevented an ISO-dependent increase in Ca^2+^ spark frequency and amplitude (Figure 4), which is known to be induced by CaMKIIδ mediated phosphorylation of RyR2 ^4^. However, we cannot rule out attenuation of Ca^2+^ cycling of other Ca^2+^ handling proteins, such as SERCA^37^ or LTCC^38^ by direct *S*-nitrosylation.

Interestingly, our novel CaMKIIδ-C273S animals had increased expression of SERCA compared to WT littermates (Figure 3), consistent with the C273S mutant mice undergoing compensation to increase SR Ca^2+^ uptake, potentially in response to increased baseline CaMKIIδ activation and enhanced calcium leak during diastole. Further work needs to be done to confirm a role for *S*-nitrosylation of CaMKIIδ as the key protein mediating these effects on calcium flux. Moreover, we acknowledge that CaMKII is subject to oxidation ^36^, and new evidence suggests that Cys residues on CaMKIIδ may be targets for oxidative stress ^39,40^. Moreover, conditions of oxidative stress that would be conducive for CaMKIIδ activation would also alter both the mechanism and rate of thiol nitrosylation^41^. Therefore, our generation of knock-in mice that either lack the Cys-273 or Cys-290 sites will be valuable tools to further investigate the effect of NO and ROS-mediated CaMKIIδ activity on cardiac function in both normal and pathological settings ^42^.

NO donors (e.g. nitroglycerin, SNP) used for their vasodilatory actions have been administered clinically for almost a century, initially for angina pectoris and later for myocardial infarction ^7^. NO donors are considered cardioprotective due in part to the ability of NO donors to mimic the benefits of ischemic pre-conditioning ^43^. However, we found GSNO or SNP pre-treatment suppressed ISO-induced cardiac arrhythmias. When the order of treatment was reversed in the isolated hearts, we found that NO donors administered after ISO could enhance the number of arrhythmias, suggesting that NO donors should be used with caution when β-AR stress is elevated. Conversely, our data demonstrate that a well-timed acute dose or low-dose chronic treatment with an NO donor (potentially paired with a CaMKII inhibitor) could be a powerful anti-arrhythmic strategy in the clinic. Importantly, our data demonstrate that the effects of NO donors on ISO-induced Ca^2+^ sparks persist even in the presence of ODQ, a soluble guanylyl cyclase inhibitor. Guanylyl cyclase is known to play a role in mediating the inotropic effects of NO in myocytes^44^, and our findings suggest that CaMKII nitrosylation works independently of guanylyl cyclase, though the two pathways may be complimentary in modulating myocytes function. These findings present new evidence for dual and opposing effects of NO signaling in the whole heart with regard to triggered arrhythmias.

## Author contributions

Experiments were performed in the laboratory of J.R.E, D.M.B., and M.J.K. within the Department of Physiology at the University of Otago, the Department of Pharmacology at the University of California, Davis, and the Department of Environmental Health and Engineering at Johns Hopkins Bloomberg School of Public Health, respectively. J.R.E. and D.M.B. are responsible for the conception of the work and interpretation of the results. A.S.P., E.A., L.P.I.W, C.A., O.V.E., R.P. and R.S.W. contributed to study design, performed and analysed experiments, prepared figures and drafted the manuscript. J.H.B. contributed to the interpretation of the work and revised the manuscript. All authors contributed to critical revisions of the manuscript.

## Acknowledgements

We thank Dr. Julie Bossuyt for advice during the course of this project.

## Supplemental Material

**Supplementary Table 1.**
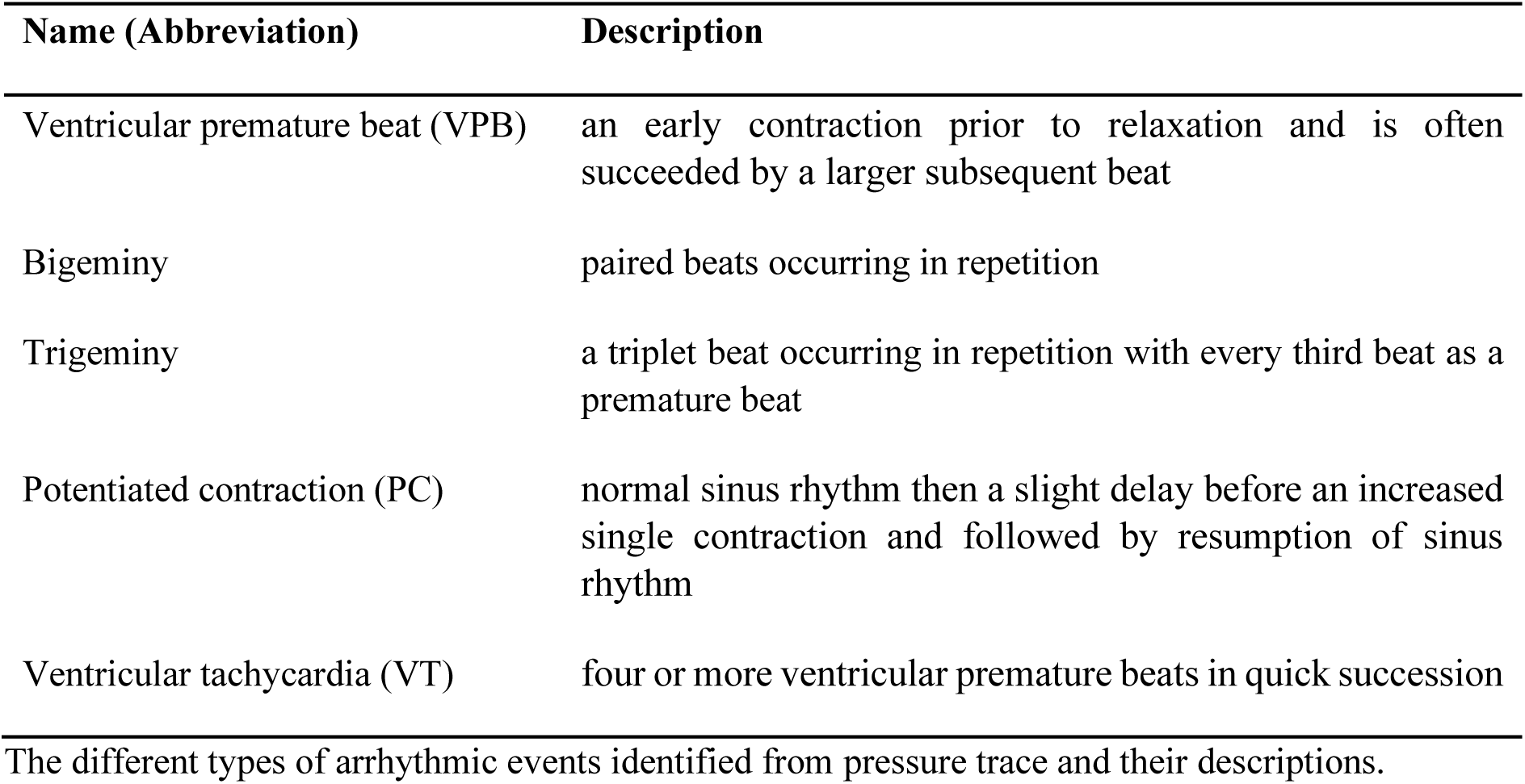
Arrhythmic events. Name (Abbreviation) Description.

**Supplementary Table 2.** Arrhythmia Score Classification. Arrhythmia Score Characteristics.

| Arrhythmia Score | Characteristics |
| --- | --- |
| 0 | 0 – 5 VPBs (no arrhythmias or isolated VPBs) |
| 1 | 6 – 20 VPBs (very occasional bigeminy and trigeminy) |
| 2 | 21 – 100 VPBs (frequent bigeminy, trigemini and PCs) |
| 3 | > 100 VPBs or 1 – 2 episodes of VT |
| 4 | > 3 VT |
| 5 | Ventricular fibrillation or death |

A 5-point arrhythmia score classification to determine severity of arrhythmias observed in the isolated perfused mouse hearts. VPB = ventricular premature beats, PC = potentiased contraction, VT = ventricular tachycardia.

**Supplementary Figure 1.**
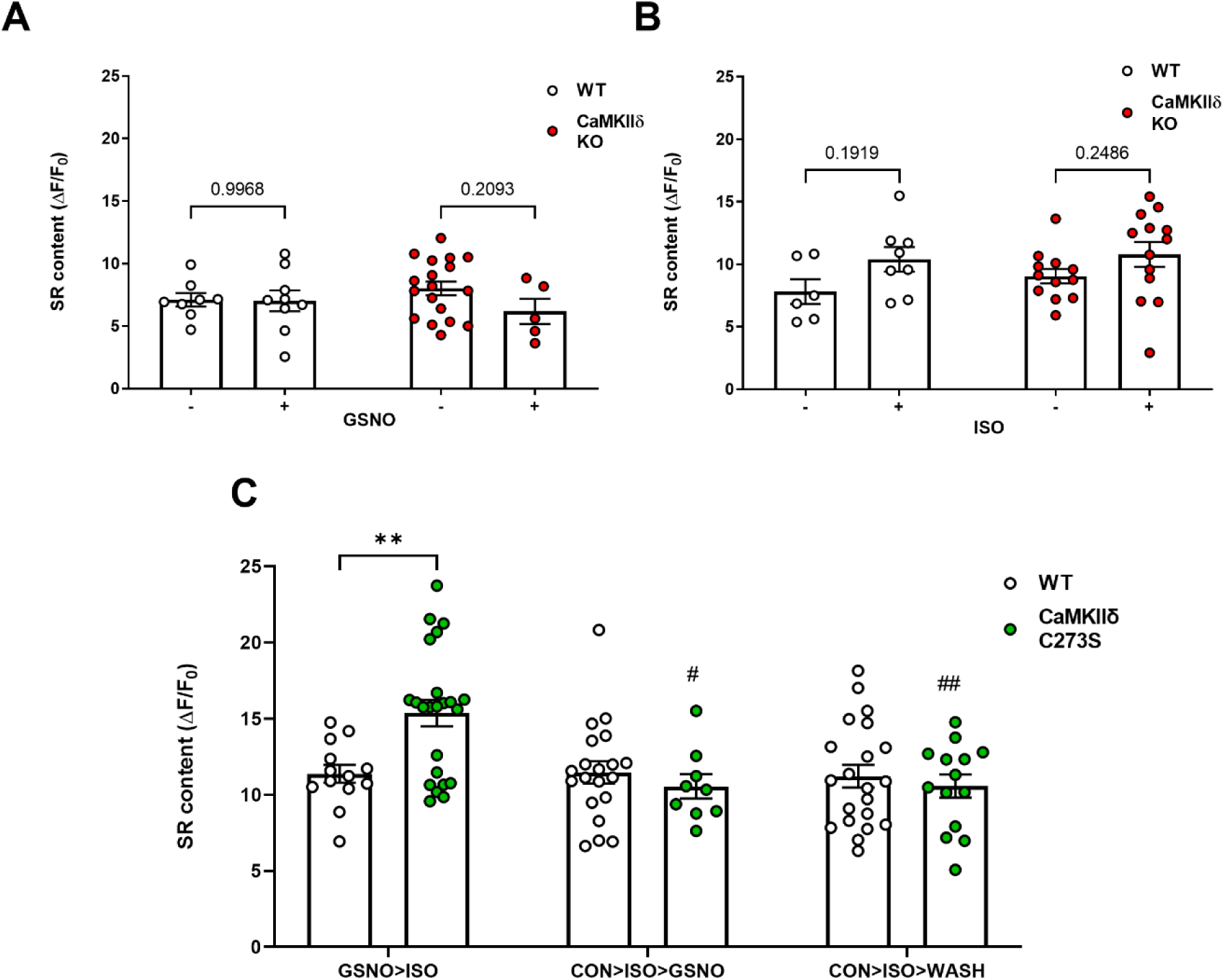
SR Ca^2+^ content. At the end of each experimental protocol, a rapid application of caffeine was used to measure SR Ca^2+^ content. Ca^2+^ SR content for WT and CaMKIIδ-KO exposed to 150 μM GSNO (**A**) or 100 nM ISO (**B**). Time controls were also performed for the WT and CaMKIIδ-KO cardiomyocytes and were not exposed to iso or GSNO. (**C**) SR Ca^2+^ content for WT and CaMKIIδ-C273S cardiomyocytes exposed to ISO after GSNO pre-treatment (GSNO>ISO; WT: n=13 cells, N=6 hearts; CaMKIIδ C273S: n=23 cells, N=8 hearts), or GSNO following ISO (CON>ISO>GSNO; WT: n=21 cells, N=9 hearts; CaMKIIδ C273S: n=9 cells, N=5 hearts) or following washout of ISO with control buffer (CON>ISO>WASH; WT: n=21 cells, N=10 hearts; CaMKIIδ C273S: n=14 cells, N=8 hearts). # p=0.001 compared to CaMKIIδ C273S GSNO>ISO, ## p=0.0001 compared to CaMKIIδ C273S GSNO>ISO.

**Supplementary Figure 2.**
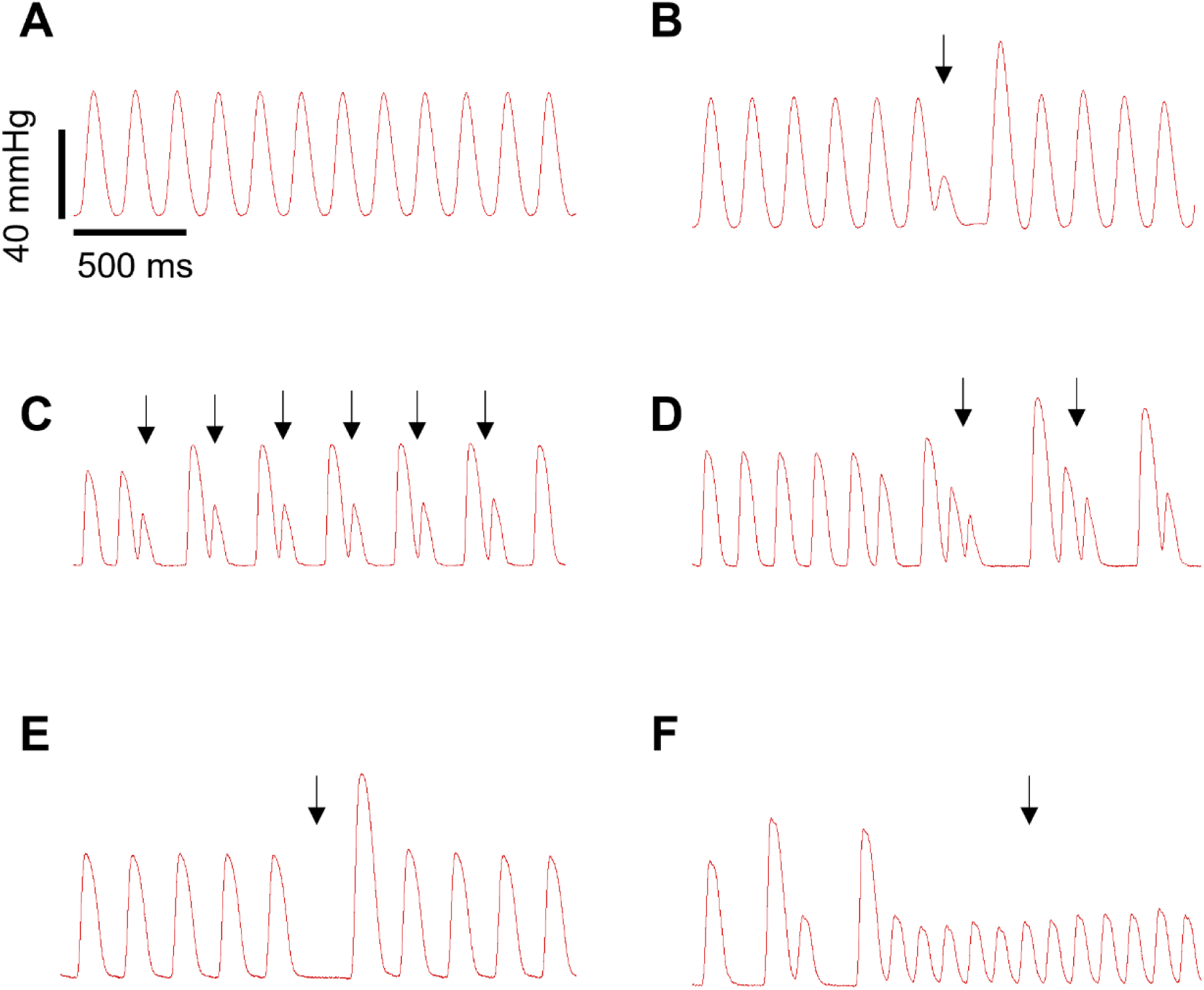
Arrhythmic events obtained from pressure trace. (**A**) sinus rhythm (**B**) ventricular premature beats (**C**) bigeminy (**D**) trigemini (**E**) potentiated contraction, and (**F**) ventricular tachycardia.

